# Integrated *in vivo* functional screens and multi-omics analyses identify α-2,3-sialylation as essential for melanoma maintenance

**DOI:** 10.1101/2024.03.08.584072

**Authors:** Praveen Agrawal, Shuhui Chen, Ana de Pablos, Faezeh Jame-Chenarboo, Eleazar Miera Saenz de Vega, Farbod Darvishian, Iman Osman, Amaia Lujambio, Lara K. Mahal, Eva Hernando

## Abstract

Glycosylation is a hallmark of cancer biology, and altered glycosylation influences multiple facets of melanoma growth and progression. To identify glycosyltransferases, glycans, and glycoproteins essential for melanoma maintenance, we conducted an *in vivo* growth screen with a pooled shRNA library of glycosyltransferases, lectin microarray profiling of benign nevi and melanoma patient samples, and mass spectrometry-based glycoproteomics. We found that α-2,3 sialyltransferases ST3GAL1 and ST3GAL2 and corresponding α-2,3-linked sialosides are upregulated in melanoma compared to nevi and are essential for melanoma growth *in vivo* and *in vitro*. Glycoproteomics revealed that glycoprotein targets of ST3GAL1 and ST3GAL2 are enriched in transmembrane proteins involved in growth signaling, including the amino acid transporter Solute Carrier Family 3 Member 2 (SLC3A2/CD98hc). CD98hc suppression mimicked the effect of ST3GAL1 and ST3GAL2 silencing, inhibiting melanoma cell proliferation. We found that both CD98hc protein stability and its pro-survival effect in melanoma are dependent upon α-2,3 sialylation mediated by ST3GAL1 and ST3GAL2. In summary, our studies reveal that α-2,3-sialosides functionally contribute to melanoma maintenance, supporting ST3GAL1 and ST3GAL2 as novel therapeutic targets in these tumors.

## Introduction

Melanoma is the most aggressive skin cancer, the incidence of which has increased rapidly over the past decades. Early staged disease can be cured by surgery; however, despite the success of targeted and immune therapies, more than 50% of patients with melanoma metastasis still succumb to their disease. Thus, identifying novel drivers of melanoma progression and maintenance is essential to developing more effective treatments for advanced disease. Aberrant epigenetic, post-transcriptional, and post-translational programs such as glycosylation may provide unique intervention points. Melanoma originates from pigment-producing melanocytes, neural crest-derived cells found in the skin, eye, and other tissues throughout the body^1, 2^. Mutations in BRAF initiate the formation of nevi, benign melanocytic lesions. While most nevi have very low levels of proliferation, histopathological data shows that approximately one-third of melanomas arise from melanocytic nevi ^3^. Transformation of nevi to melanoma in situ occurs through crosstalk between mutated cancer cells and their immediate microenvironment ^4, 5^. As primary tumor cells override tumor-suppressing barriers via oncogenic adaptations, invasive melanoma cells grow into the dermis and access systemic circulation ^1^. Cells surviving in circulation can extravasate, survive and colonize regional and distant organs ^2, 6^.

Altered glycosylation is a hallmark of cancer biology ^7–9^ that influences multiple facets of tumor growth and progression^10^. Recent findings of altered glycosylation in melanoma include upregulation of branching and bisecting *N-*linked glycans and truncated *O-*linked glycans contributing to the invasive and metastatic properties of melanoma ^11–14^. In our previous work, we analyzed patient samples and identified core fucosylation, incorporated by α-1,6 fucosyltransferase FUT8, as a driver of melanoma metastasis ^15^. However, current knowledge of glycans and their role in promoting melanoma growth and maintenance remains incomplete.

Herein, we conducted the first *in vivo* functional screen of glycosyltransferases in melanoma using a pooled library of doxycycline-inducible shRNAs. This analysis revealed a consistent depletion of melanoma cells carrying shRNAs targeting the family of α-2,3-sialyltransferases during *in vivo* growth. Concordantly, lectin array profiling of nevi vs. melanoma patient samples revealed enhanced α-2,3-sialylation, and *in silico* analyses of clinical transcriptomic datasets identified *ST3GAL1* and *ST3GAL2* as consistently upregulated in melanoma vs. nevi. Lectin pull-downs followed by mass-spectrometry analysis identified sialylated proteins responsible for promoting melanoma growth, including the amino acid transporter Solute Carrier Family 3 Member 2 (SLC3A2, CD98hc). Overall, our work sheds light on the role of glycosylation in melanoma biology and opens a novel path to developing glycan-based therapeutics to treat melanoma patients.

## Results

### Integrated *in vivo* shRNA functional screen and multi-omics analyses identify glycogenes and glycans essential for melanoma growth

To identify glycogenes and their respective glycan epitopes required for melanoma cell survival, we utilized a multi-omics approach, integrating a functional *in vivo* growth screen of glycogenes with high throughput glycomic profiling of clinical melanoma samples (Fig. 1A). To identify glycogenes essential for melanoma growth, we designed a pooled library of doxycycline-inducible small hairpin RNA (shRNA) targeting glycan biosynthetic enzymes^16^. Our retrovirus-based shRNA library targets 199 glycosyltransferase genes, representing the majority of the 214 annotated in the human genome^17^ and contains six shRNAs/gene. These pooled libraries were divided into sub-libraries of 100 shRNAs to get sufficient coverage during next-generation sequencing (NGS) of xenograft tumors (Fig. 1B). Sub-libraries were used to transduce MeWo melanoma cells implanted subcutaneously in immunocompromised mice. To identify glycogenes essential for melanoma growth *in vivo*, we performed NGS of tumor cells before implantation and at 5 weeks post-injection. Our analysis revealed multiple consistently depleted shRNA (average fold change <0.6, t-test p<0.05, at least 3/6 shRNA read counts in tumors vs. baseline, Fig. S1A-B). The following glycogenes were found to be depleted: *B3GAT1*, *B3GNT4*, *B3GNT5*, *GALNT2*, *GCNT2*, *HS3ST6*, *OGT*, *RFNG*, *RPN2*, *ST3GAL1*, *ST3GAL2*, *ST3GAL5*, *ST3GAL6*, *ST6GALNAC2*, *ST8SIA2*, *UGT8* and *XYLT1* (Fig. 1C, Fig S1B). OGT and RPN2 are known to be essential for cell viability ^18–20^, confirming the ability of our growth screen to identify essential genes. Of note, multiple members of the α−2,3 sialyltransferase family (*ST3GAL1/2/5/6*) showed depleted shRNAs in tumors, indicating a potential role in maintaining melanoma growth (Fig. 1C, Fig. S1B).

**Fig. 1.**
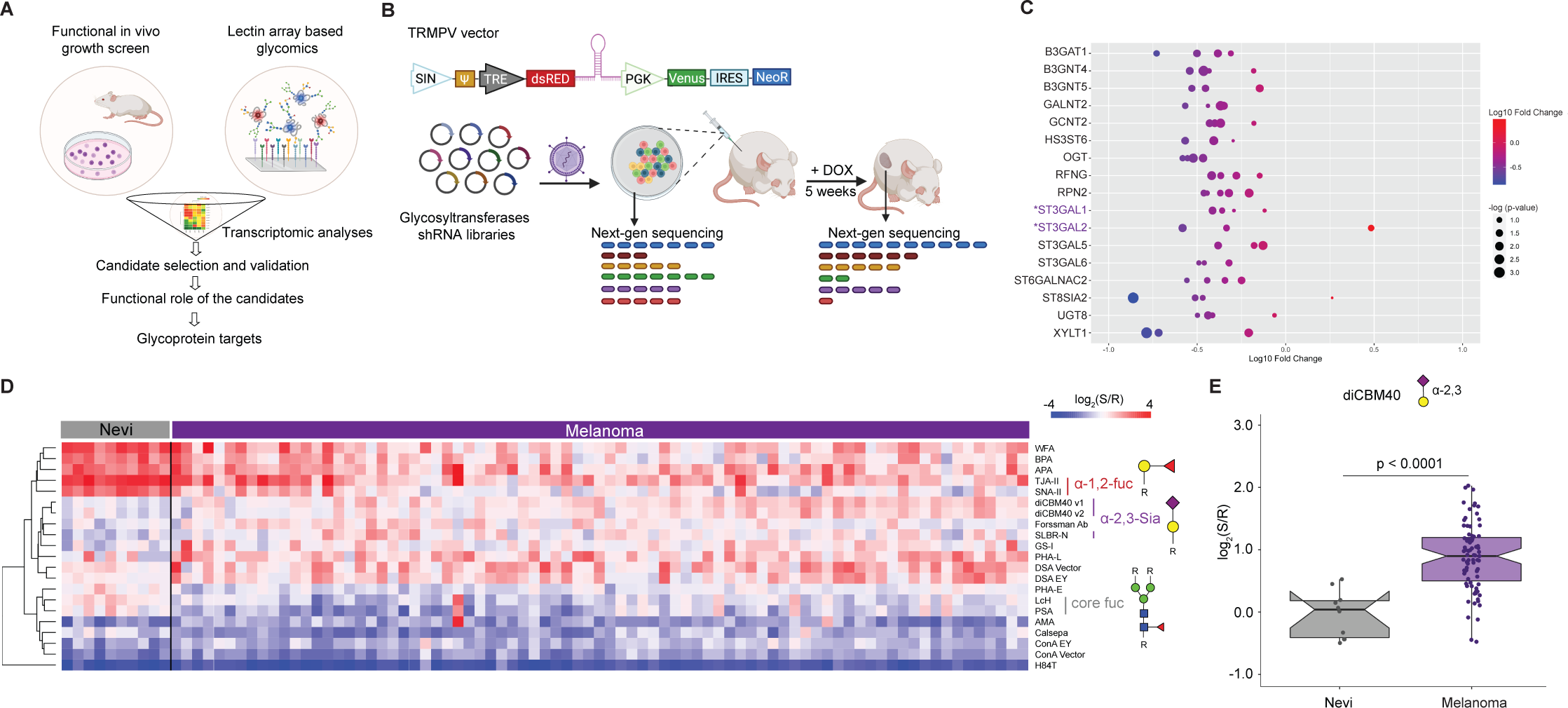
*In vivo* functional genetic screen and multi-omics approach identify essential glycogenes and glycans for melanoma growth. **(A)** Schematic illustration of our approach to identifying glycosylation enzymes involved in melanoma growth and their targets. **(B)** Schematic representation of dual-color TRINE vector that enables Tet-regulated shRNA expression to suppress glycosyltransferases involved in cell proliferation and survival. The *in vivo* growth screen schema is also presented. **(C)** Bubble plot representing log 10-fold change of the depleted or enriched shRNA in the MeWo cells transduced with glycosyltransferase shRNA libraries in tumors (5 weeks post-injection) relative to baseline (before injection). Glycosyltransferases corresponding to α-2,3 sialylation are highlighted in purple. **(D)** Heatmap of ratiometric lectin microarray data for nevi (n=10) and melanoma FFPE tissues (n=79); Only lectins showing significant differences between the 2 groups are shown (student’s t-test (two-tailed), *p* < 0.05). Pink, log2(S/R) > log2(Smedian/Rmedian); blue, log2(Smedian/Rmedian) > log2(S/R). Lectins corresponding to α-2,3 sialosides are highlighted in purple. The complete heatmap is given in Fig. S2A. (**E**) Whisker plot showing significantly increased diCBM40 binding in melanoma compared to nevi. Significance was determined using Wilcoxon’s t-test.

To complement our functional *in vivo* screens and identify glycosylation signatures associated to melanoma growth and progression, we analyzed melanocytic nevi (n=18) and melanoma (primary, n=10; metastatic, n=61) FFPE patient samples using lectin microarrays. Melanocytic nevi are benign lesions unlikely to progress to melanoma; however, accumulating evidence suggests they are melanoma precursors ^2, 3, 21^. The inclusion of nevi thus allowed the identification of glycan changes happening during melanomagenesis. Formalin-fixed paraffin-embedded (FFPE) melanoma tissue samples were processed and analyzed using our dual-color lectin microarray technology ^15, 22^. Lectin microarrays utilize carbohydrate-binding proteins with well-defined specificities ^23^ to detect glycan changes at different stages of melanoma progression. Heatmaps (Figures 1D and S2A) show differences in lectin binding between nevi and melanomas (including primary, lymph node metastases, subcutaneous metastases, and brain metastases). In keeping with our earlier work, we see a loss of α-1,2 fucose (lectins: TJA-II, SNA-II) ^24^ in primary melanoma compared to nevi that is even more striking in the metastasis (Fig. 1D). Interestingly, core fucose (lectins: LcH, PSA) is high in nevi and lower in primary melanoma but then appears to be regained in metastatic melanoma where we have shown it contributes to invasion ^15^. We also observed a significant increase in binding to lectins diCBM40 ^25^ and SLBR-N ^26^, which recognize α-2,3-sialosides, in both primary and metastatic melanoma compared to nevi (Fig. 1D-E and S2B). Combined with the results of our functional selection experiment, our data points to a role for members of the sialyltransferase family and α-2,3-sialosides in melanoma growth and survival.

### ST3GAL1 and ST3GAL2 display higher transcriptional and protein levels in melanoma compared to nevi

Our *in vivo* growth screens and glycomic analysis of benign and melanoma patient samples revealed multiple candidate regulators of melanoma growth. To narrow the list, we further filtered candidates by transcriptomics analysis of melanoma patient cohorts with the expectation that essential genes would show transcriptional upregulation in melanoma (Fig. 1A). We observed that most of the glycogene candidates from our *in vivo* growth screens did not show significant transcript differences in benign vs. melanoma samples (Fig. 1A, Fig. S3A-C). While *ST3GAL1* and *ST3GAL2* transcripts were significantly upregulated in melanoma compared to benign nevi (Fig. 2B) in multiple datasets. These analyses suggest that ST3GAL1 and ST3GAL2 upregulation is associated with neoplastic transformation and could contribute to melanomagenesis. ST3GAL1 and ST3GAL2 are sialyltransferases that catalyze the incorporation of α-2,3 sialic acids onto Galβ1,3GalNAc acceptors on glycoproteins and glycolipids (Fig. 2A) ^27^. ST3GAL1 transfers sialic acid onto *O*-linked glycans (e.g., mucins) ^28^, while ST3GAL2 can biosynthesize both *O*-glycans and glycolipids^29, 30^. Immunohistochemistry (IHC) analysis of an independent cohort of nevi and melanoma tissue microarray confirmed higher ST3GAL1 and ST3GAL2 protein in melanoma as demonstrated by perinuclear granular staining (Fig. 2C, Fig. S4A). In concordance with this, we observed significantly increased staining by diCBM40, a lectin that binds α-2,3-linked sialosides, in melanoma vs. nevi ^25^ (p<0.001; Fig. 2D). We confirmed the specificity of diCBM40 with α-2,3-specific neuraminidase (NEB, P0743), which diminished the staining from this lectin (Fig. S4B). Our analyses suggest that transcriptional upregulation of ST3GAL1 and ST3GAL2 in melanoma results in increased sialyltransferase activity and higher levels of α-2,3-sialylated proteins. We postulated that ST3GAL1 and ST3GAL2 upregulation in melanoma is pro-tumorigenic and/or positively selected during tumorigenesis to support survival and growth during transformation.

**Fig. 2.**
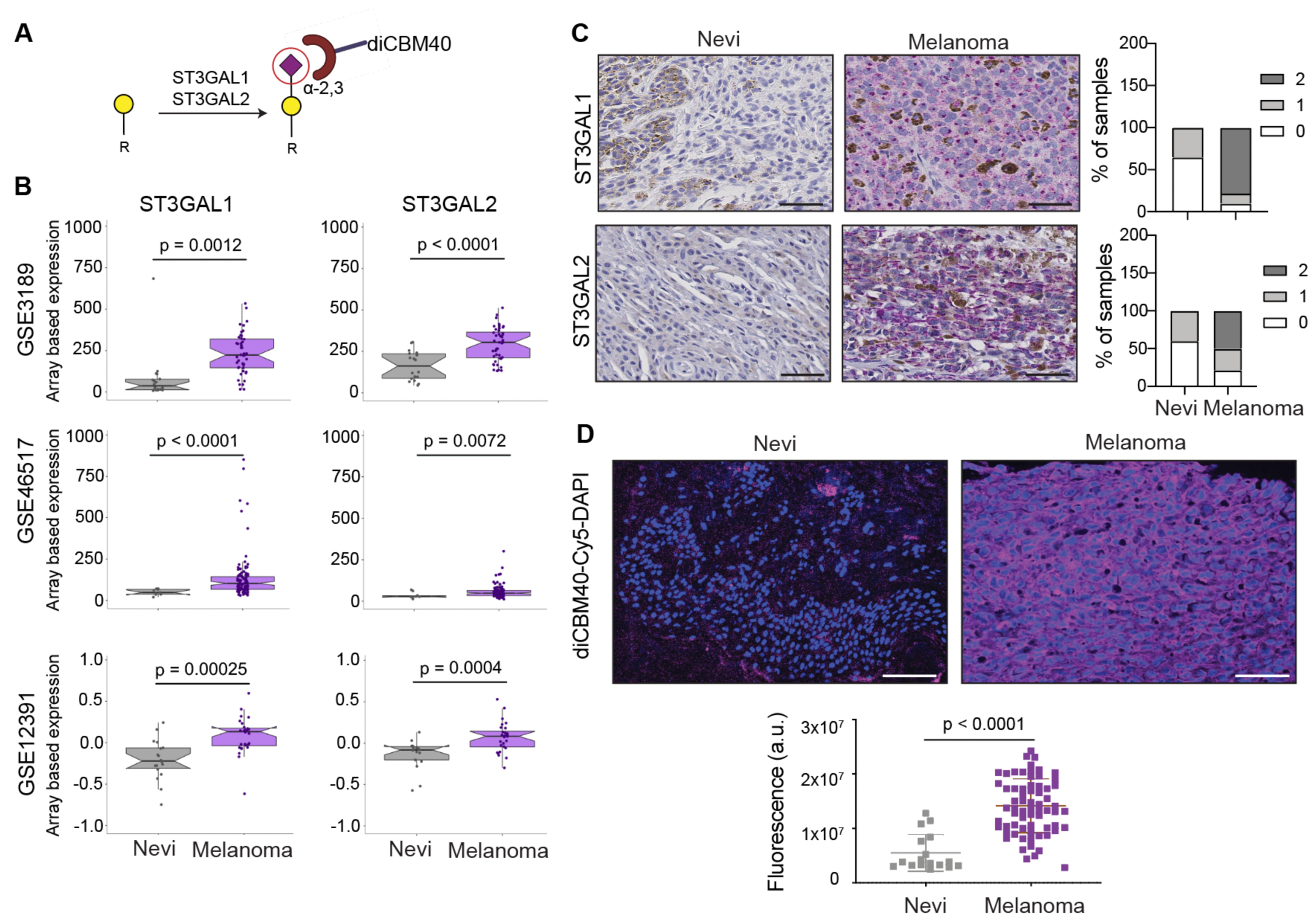
ST3GAL1, ST3GAL2, and α-2,3-sialosides are upregulated in melanoma relative to nevi. **(A)** α-2,3 sialylated glycans generated by ST3GAL1 and ST3GAL2 and recognized by diCBM40 lectin. **(B)** Whisker plot illustrating significant upregulation of *ST3GAL1* and S*T3GAL2* mRNA expression in melanoma samples compared to nevi in multiple datasets: GSE3189^81^, GSE46517^82^, GSE12391 ^83^. Two-tailed unpaired t-test. **(C)** Representative images of IHC staining with ST3GAL1 and ST3GAL2 antibodies in 15 nevi and 50 melanoma samples show a perinuclear staining pattern (Fast red counterstaining). IHC score was calculated by combining the signal intensity and percentage of positive cells within the section. The histogram shows the distribution of ST3GAL1 and ST3GAL2 IHC scores in nevi and melanoma samples. Scale bar, 10 µm. **(D)** Representative images of diCBM40 lectin fluorescence microscopy of nevi and melanoma FFPE tissues (n = 17 for nevi and 68 for melanomas). diCBM40-Alexa 647 (magenta) and DAPI-stained sections of TMA. Scale bar, 100 µm. Dot plots represent the average fluorescence intensity of five fields per image, two-tailed unpaired t-test.

### ST3GAL1 and ST3GAL2 are essential for melanoma proliferation *in vitro*

To further investigate the role of α-2,3-sialylation in growth, we silenced ST3GAL1 or ST3GAL2 in a human metastatic melanoma cell line 131/4-5B1 (hereafter, 5B1) ^31^ and a patient-derived short-term culture, 12-273BM ^32^. In brief, 5B1 and 12-273BM cells were stably transduced with lentivirus carrying one of two independent short hairpin RNAs (shRNAs) targeting *ST3GAL1* (shA or shB) or *ST3GAL2* (shC or shD) or a non-targeting, scrambled control (shSCR). We confirmed ST3GAL1/2 knockdown by qRT-PCR and Western blot (WB) (Fig. 3A-B, Figure S5A-B). Silencing of *ST3GAL1* or *ST3GAL2* substantially decreased binding by diCBM40, implying a concomitant reduction in α-2,3-linked sialosides (Fig. S5C). In both cases, silencing these genes also substantially decreased melanoma cell proliferation (Fig. 3C). No impact on cell growth was observed in normal melanocytes (NHEM) or in non-melanoma cell lines derived from the human embryonic kidney (HEK293) and lung adenocarcinoma (A549, Fig. S5D-E). Knockout of these sialyltransferases in mouse models have been generated by others and are viable ^33, 34^, arguing that the anti-proliferative effect observed is not general to all cell types and may be specific to melanoma. To further characterize this growth defect, we investigated the effects of *ST3GAL1 and ST3GAL2* knockdown on apoptosis. Flow cytometry analysis for Annexin V- and PI positive cells (Fig. 3D), together with WB for cleaved PARP and caspase-3 (Fig. 3E), all classical apoptosis markers, demonstrate that melanoma cells in which *ST3GAL1* or *ST3GAL2* are knocked down indeed undergo apoptosis. Moreover, simultaneous incubation with pan-caspase inhibitor Q-VD-OPh impaired melanoma cell death (Fig. S5F), further supporting caspase-dependent apoptosis in *ST3GAL2* silenced cells.

**Fig. 3.**
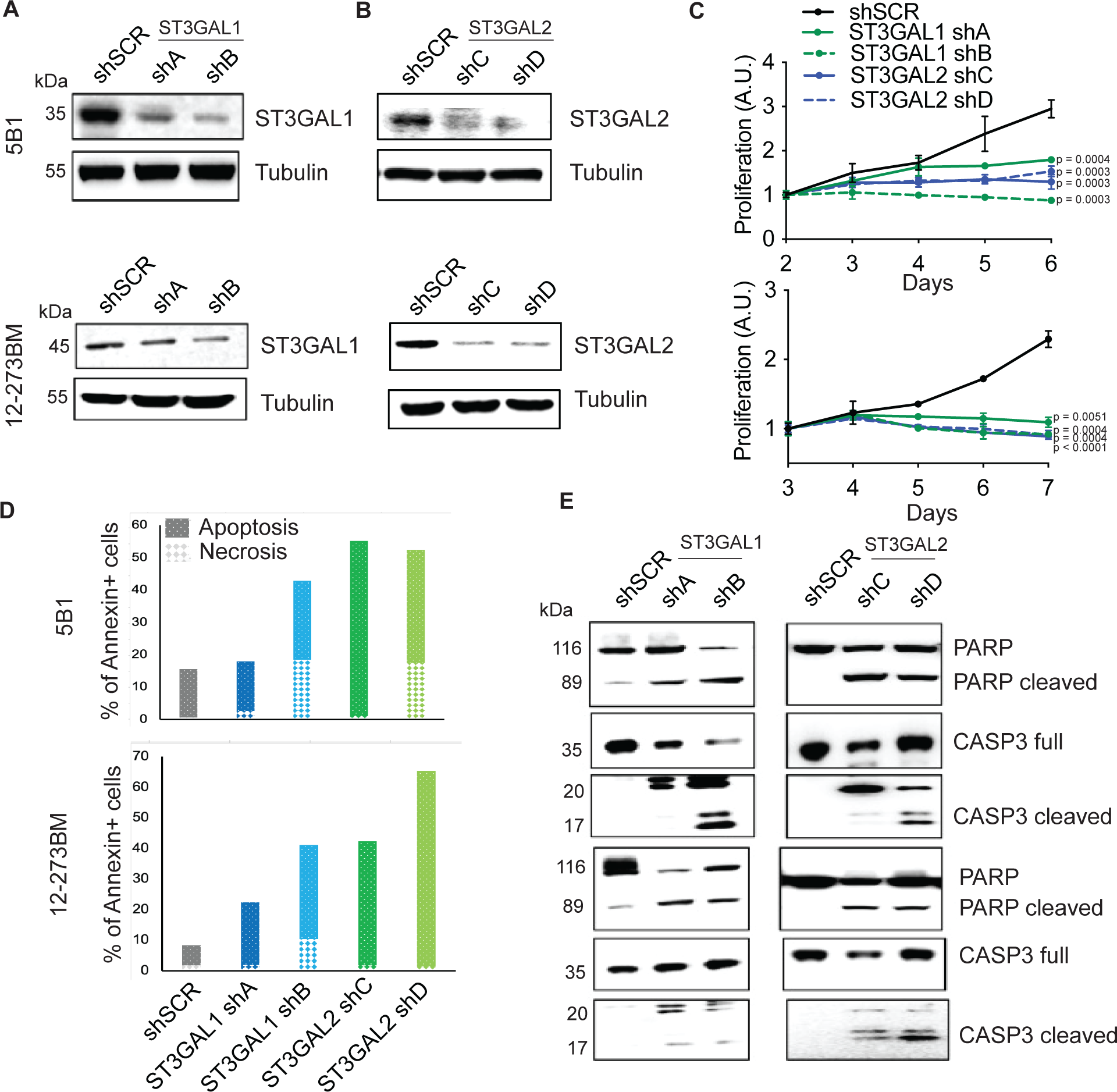
ST3GAL1 and ST3GAL2 are essential for melanoma proliferation *in vitro*. ST3GAL1 **(A)** and ST3GAL2 **(B)** protein levels in 5B1 and 12-273BM cells stably transduced with non-targeting scrambled control shRNA (shSCR), *ST3GAL1* shRNAs (shA and shB) and *ST3GAL2* shRNAs (shC and shD) were assessed by Western blotting. Western blot images are representative of three independent experiments. **(C)** Relative growth curves of 5B1 and 12-273BM cells stably transduced with non-targeting scrambled control shRNA (shSCR), *ST3GAL1* shRNAs (shA and shB), and *ST3GAL2* shRNAs (shC and shD). The data shown are representative of three independent experiments. Two-tailed unpaired t-tests and p-values are shown in the figures. **(D)** The percentage of melanoma cells positive for Annexin V only (early apoptosis) or PI only (necrosis), Experiment was performed in duplicates. **(E)** Representative Western blots for caspase 3 and PARP on lysates from 5B1 and 12-273BM cells with shRNA against ST3GAL1 and ST3GAL2.

### Identification of α-2,3-sialylated glycoproteins in melanoma reveals novel regulators of viability and maintenance

To identify α-2,3-sialylated glycoproteins that could mediate the role of ST3GAL1 and ST3GAL2 in melanoma proliferation, we performed proteomic analysis of α-2,3-sialylated membrane proteins. We collected membrane extracts from three melanoma cell lines, 5B1 (BRAF mutant), 12-273BM (NRAS mutant), and MeWo (NF1 mutant/BRAF wild type/NRAS wild type), which represent the major melanoma genotypes, to identify shared sialylated glycoproteins. The α-2,3-sialylated glycoproteins were enriched using lectin chromatography with MAA (Fig. 4A). We confirmed α-2,3-sialylated glycoprotein enrichment by silver staining and MAA lectin blot (Fig. S6), and samples were analyzed by liquid chromatography coupled with mass spectrometry (LC-MS). As expected, less MAA-bound protein was observed from ST3GAL1, and ST3GAL2 silenced cells (Fig. 4B). Mass spectrometry data were processed by median filtering. Contaminant proteins such as ribosomal proteins, tubulins, and heat shock proteins are known to be non-glycosylated and were removed from the analysis. We identified 85, 78, and 75 candidate proteins in the MAA enrichment of 12-273BM, 5B1, and MeWo, respectively, and 44 out of the total proteins were common to all (Fig. 4C). Gene ontology analysis (DAVID) of shared sialylated proteins revealed enrichment in growth, locomotion, and adhesion factors (Fig. 4D). Proteins bound by MAA within the cell growth category include Transferrin receptor 1 (TFR1), Solute carrier family three member 2 SLC3A2 (also known as CD98 heavy chain (hc)), Apolipoprotein E (ApoE), Solute carrier family one member 4 (SLC1A4), Serpin Family H Member 1 (SERPINH1), and Galectin-3 binding protein (LGALS3BP) (Table S1). We investigated the potential association of these genes with the survival of melanoma cells using the Dependency Map (DepMap) ^35–37^, which catalogs genetic dependencies in cancer cells by pooled RNAi or CRISPR screens. We observed that many melanoma cell lines and cells of other cancer types are frequently dependent on *TFR1* and *SLC3A2* for survival (Fig. S7 A-C). 32

To validate our proteomic analysis, we further examined the glycosylation state of TFR1 and CD98hc. MAA lectin enrichment followed by Western blot showed reduced TFR1 and CD98hc levels in shA (shST3GAL1) and shC (shST3GAL2) compared with shSCR-transduced 5B1 cells (Fig. 4E), consistent with lower α-2,3-sialylation on those proteins. Interestingly, silencing of ST3GAL1/2 also led to reduced total CD98hc protein levels (Fig. 4E,F). In contrast, TFR1 levels did not change in ST3GAL1-silenced cells and were found to increase in shST3GAL2-transduced cells (Fig. 4F). Overall, our analysis suggests CD98hc and TFR1 are α-2,3-sialylated by ST3GAL1 and ST3GAL2 and that CD98hc levels may be influenced by sialylation.

**Fig. 4.**
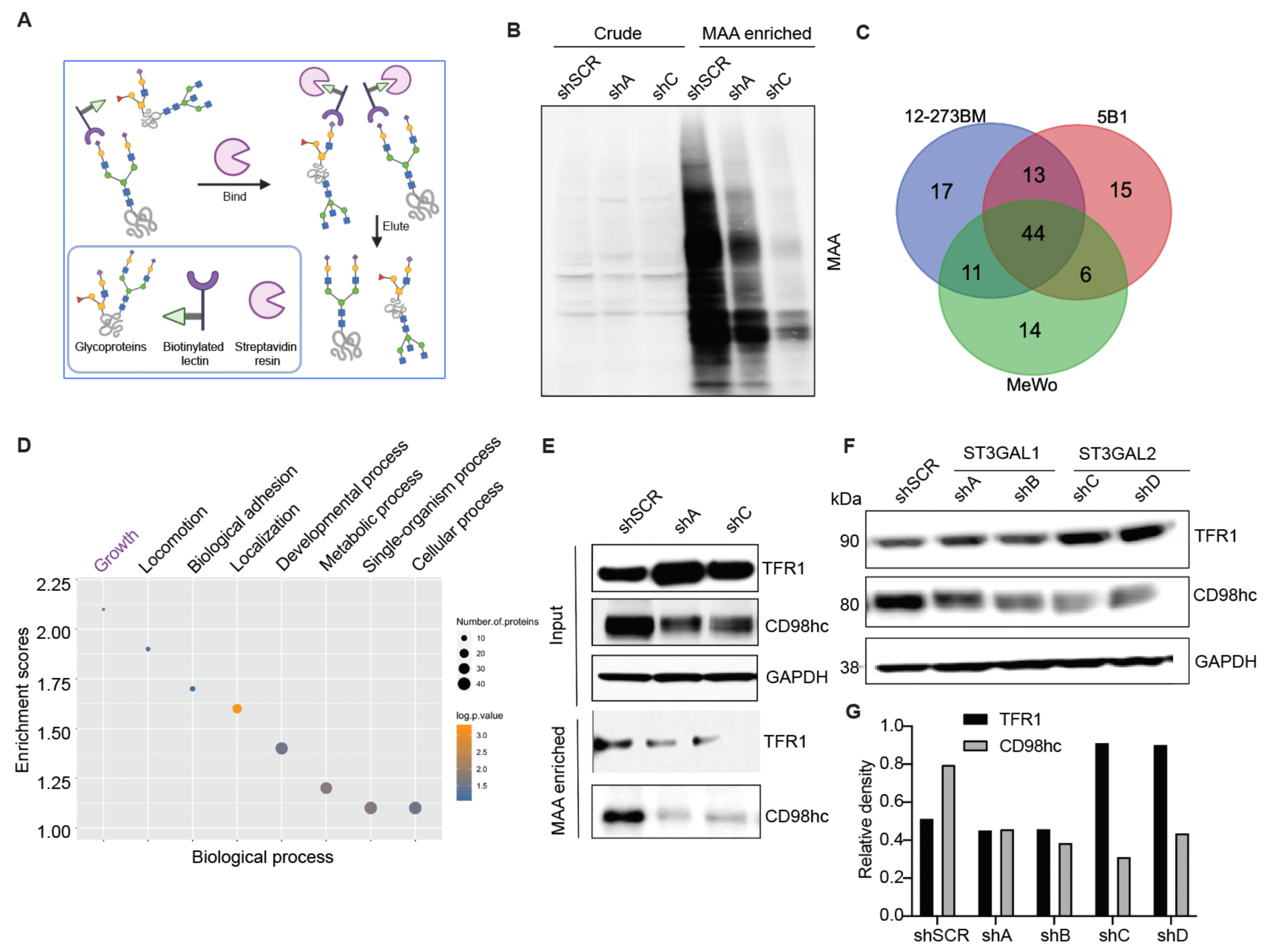
Identification of α-2,3-sialylated glycoproteins in melanoma reveals regulators of melanoma growth. **(A)** Schematic illustration of the experimental approach showing affinity enrichment of α-2,3 sialylated proteins by MAA lectin affinity chromatography. **(B)** MAA affinity chromatography of whole-cell lysates of 5B1 cells transfected with NTC or ST3GAL1/ST3GAL2 shRNA followed by lectin blot with MAA lectin. An equal concentration of crude protein and an equal volume of MAA-enriched fractions was loaded for MAA lectin blot. **(C)** A number of proteins were identified by mass spectrometry analysis of the MAA-enriched fractions from 5B1, 12-273BN, and MeWo cell lines. (**D**) Gene ontology enrichment analysis (biological processes category) of α-2,3 sialylated proteins common to the three cell lines. Also, see Table S1 **(E)** Western blot analysis with α-TFR1 or α-CD98hc antibodies of MAA-pulldown and corresponding input from lysates of 5B1 cells transfected with NTC or ST3GAL1/ST3GAL2 shRNA. **(F)** Western blot analysis of TFR1 and CD98hc in lysates from 5B1 cells transfected with NTC or ST3GAL1 or ST3GAL2 shRNAs. (**G**) Densitometric analysis of CD98hc on lysates from 5B1 cells transfected with NTC or ST3GAL1 or ST3GAL2 shRNAs. The graph is representative of three replicates. Experiments in B, E, and F were performed in triplicate, and representative images were shown.

### α-2,3-sialylation of SLC3A2 (CD98hc) is required for its stability and anti-proliferative effect

CD98 heavy chain (CD98hc), encoded by SLC3A2, consists of a transmembrane domain, a heavily glycosylated extracellular domain, and a cytoplasmic tail ^38, 39^. SLC3A2 is then disulfide-linked with the non-glycosylated light chains (CD98lc) encoded by either LAT-1 or LAT-2 ^40^. CD98 participates in multiple biological functions, including amplifying integrin signaling ^41^ and mediating amino acid transport ^42^, which ultimately impact cell proliferation and survival. Moreover, altered SLC3A2 expression has been associated with poor prognosis in several types of cancer ^43–46^. A recent study showed that SLC3A2 was highly expressed in metastatic melanoma tissue compared to benign melanocytic nevi, and higher *SLC3A2* mRNA levels were strongly correlated with lower survival rates ^47^, proposing SLC3A2 as a prognostic biomarker for melanoma. We found that silencing of CD98hc suppresses melanoma cell proliferation (Fig. 5A-B) and has no effect on other cell types, such as HEK 293T (Fig. S8), mimicking the effects of ST3GAL1 or ST3GAL2 silencing on those cells. We also found that CD98hc overexpression partially rescues the antiproliferative effect of ST3GAL1- or ST3GAL2 silencing (Fig. 5C-E; Fig. S9), suggesting that CD98hc may be a critical mediator of ST3GAL1/2 effects on melanoma growth.

**Fig. 5.**
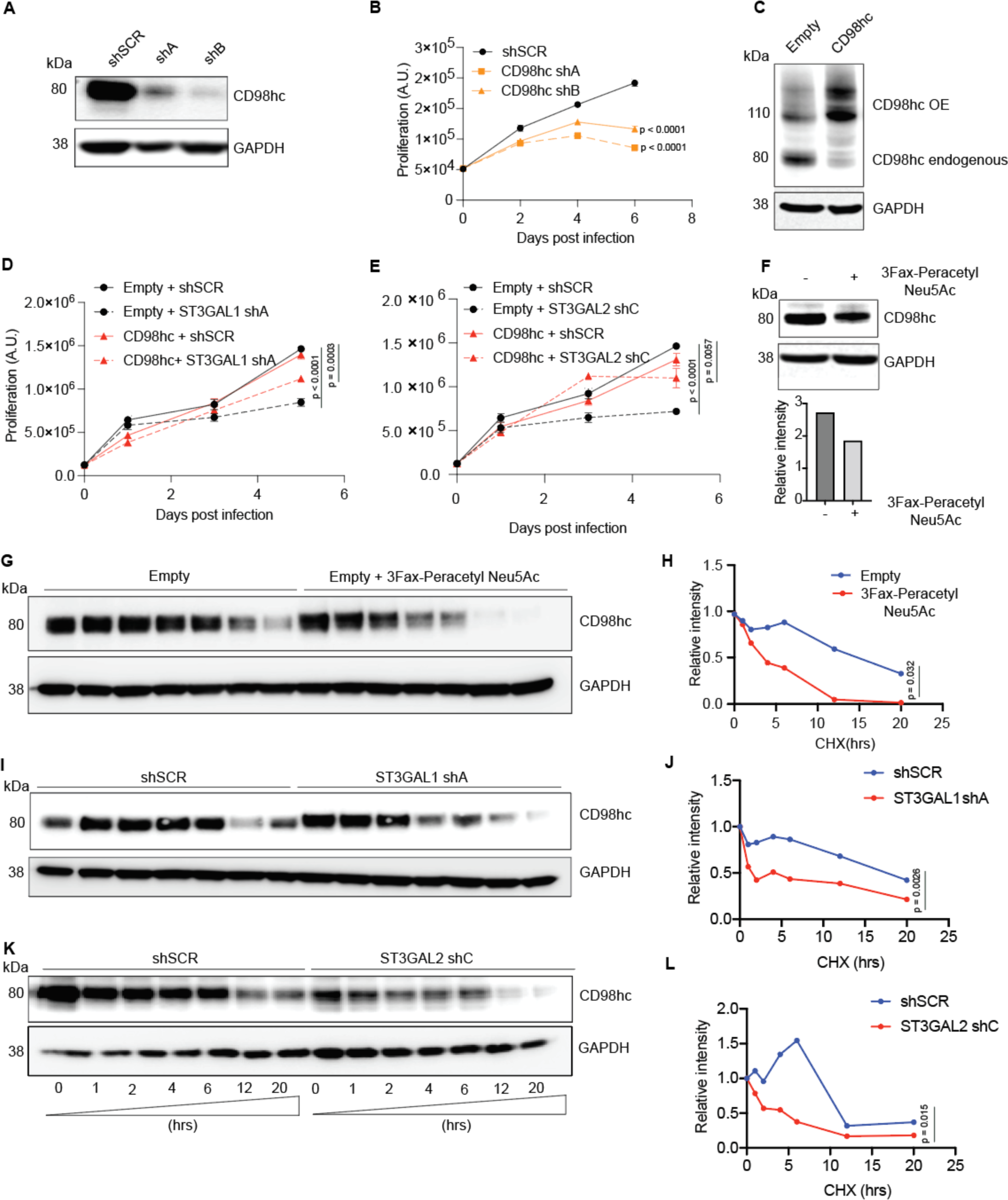
α-2,3 sialylation of SLC3A2 (CD98hc) is required for its stability and anti-proliferative effect. (**A**) Western blot for CD98hc on lysates from 5B1 cells transfected with NTC or CD98hc shRNAs. (**B**) Relative growth curves of 5B1 cells stably transduced with non-targeting control shRNA (shSCR) and CD98hc shRNAs (shA and shB). The data shown are representative of two independent experiments. Two-tailed unpaired t-tests and p-values are shown in the figures. (**C**) Western blot of CD98hc in 5B1 cells stably overexpressing CD98hc or control vector. (**D**) Cell proliferation assay on 5B1 melanoma cells stably overexpressing CD98hc or empty vector and transduced with non-targeting control shSCR or shST3GAL1. (**E**) Cell proliferation assay on 5B1 melanoma cells stably overexpressing CD98hc or empty vector and transduced with non-targeting control shSCR or shST3GAL2. (**F**) Western blot of CD98hc in 5B1 cells treated with or without 200 uM 3Fax-peracetylNeu5Ac. Densitometric analysis is shown below. (**G-H**) Western blot analysis of CD98hc protein in 5B1 cells. 3Fax-peracetylNeu5Ac treated 5B1 cells were further treated with 10 μM CHX at indicated intervals and analyzed by western blot analysis. The intensity of CD98hc protein was quantified using a densitometer. Western blot analysis of SLC3A2 protein in 5B1 cells silenced for ST3GAL1 (**I-J**) or ST3GAL2 (**K-L**) and treated with 10 μM cycloheximide (CHX) at indicated intervals and analyzed by western blot analysis. Paired t-test is shown. Western blot images are representative of 2 independent experiments.

We next sought to determine whether α-2,3-sialylation affects CD98hc protein stability. We used 3Fax-Peracetyl Neu5Ac, a cell-permeable sialic acid analog inhibitor of sialyltransferases, to assess the effect of blocking sialylation on CD98hc stability. As expected, 3Fax-Peracetyl Neu5Ac treatment reduced sialic acid-specific MAL-2 lectin binding while increasing Galβ1-3 GalNAc specific PNA lectin binding (Fig. S10A-B). Indeed P-3F_AX_-Neu5Ac treatment mimicked the effect of ST3GAL1 or ST3GAL2 silencing by reducing CD98hc levels (Fig. 4F, 5F). Moreover, in the presence of protein synthesis inhibitor cycloheximide (CHX), the turnover rate for de-sialylated CD98hc was much faster than for sialylated CD98hc (Fig. 5G-H). Similarly, ST3GAL1 or ST3GAL2 silencing resulted in more rapid CD98hc degradation in the presence of CHX (Fig. 5I-L). Collectively our data indicates that α-2,3-sialylation supports the protein stability of CD98hc, an amino acid transporter required for melanoma cell survival (Fig. 6).

**Fig. 6.**
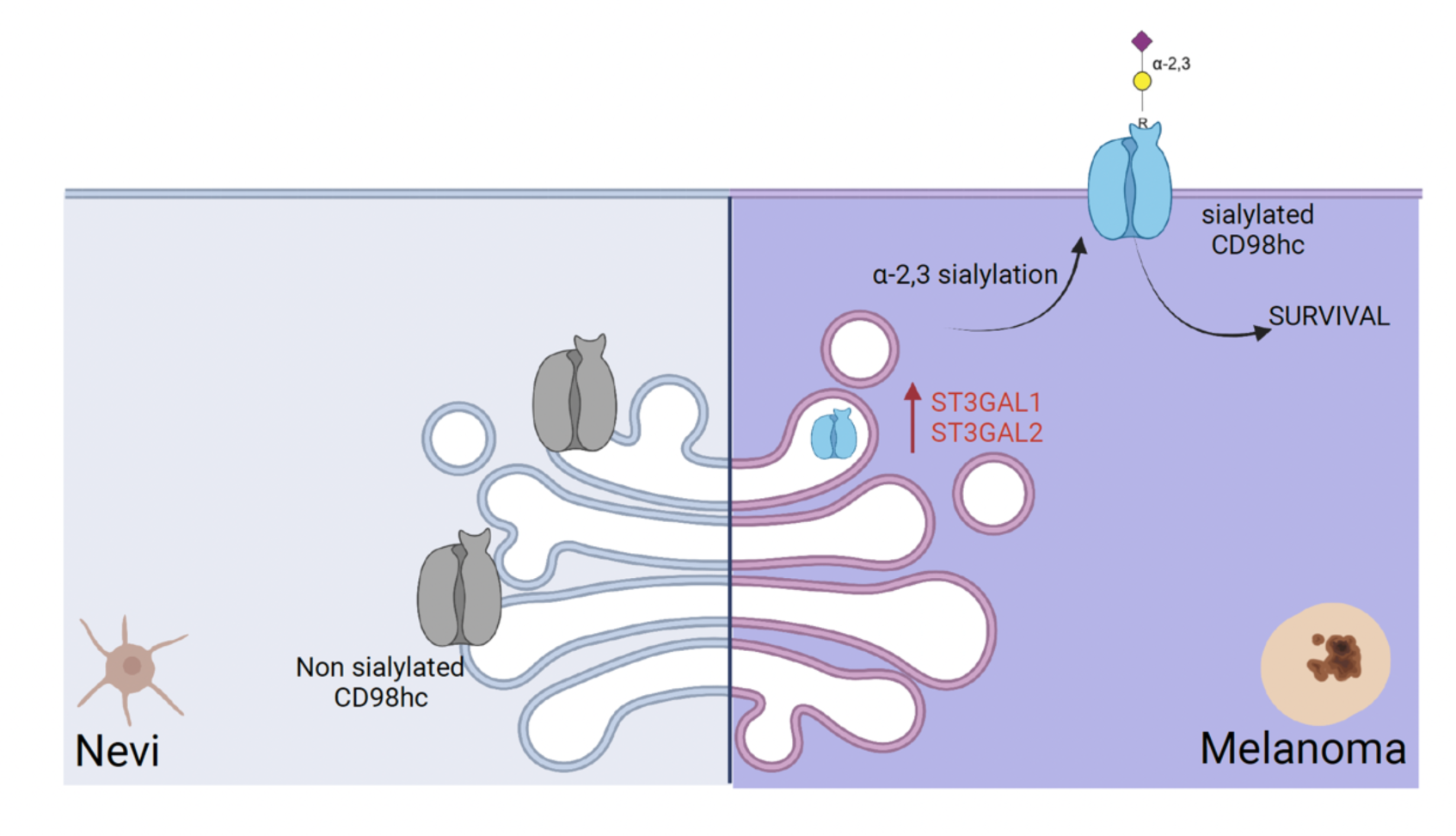
A schematic model suggests that α-2,3 sialylated transmembrane glycoprotein is required for melanoma cell survival.

## Discussion

Aberrant glycosylation influences multiple facets of malignant transformation and tumor progression^10^. In our previous work, we obtained glycan profiles of matched primary and metastatic melanomas using clinical specimens and identified core fucosylation generated by FUT8 as a molecular driver of melanoma metastasis ^15^. However, little is known about glycans’ role in melanoma genesis and maintenance. This study provides experimental and clinical evidence of the role of α-2,3 sialyltransferases in melanoma cell growth. Further, we identify α-2,3 sialylation of solute transporter protein CD98hc as a key regulator of melanoma growth.

To identify glycogenes and their glycan epitopes required for melanoma survival, we utilized a multi-omics approach, including an unbiased analysis of glycogenes using a functional *in vivo* growth screen. The decision to conduct the screen *in vivo* was based on the fact that *in vivo* models provide a more physiologic context for studying requirements for melanoma growth than *in vitro* systems. Our *in vivo* shRNA library consisting of relatively small shRNA pools (100 different shRNAs/sub-library), allowed us to examine the role of glycosyltransferases with high resolution. Supporting the value of this strategy, our screen identified glycogenes like OGT and RPN2 previously implicated in cancer cell survival ^18, 20^. In addition, our analysis identified multiple members of the sialyltransferases family ^48^, some of which are known to play an essential role in cancer cell survival, immune regulation, and angiogenesis (Fig. 1B)^49, 50^.

Most studies of glycan alterations in melanoma have been limited to *in vitro* assays on cell lines ^11, 12, 51–53^. Our parallel high-throughput lectin profiling of benign nevi and clinical melanoma samples addresses this prior limitation. We observed increasing levels of α-2,3-sialosides (diCBM40, SLBR-N) in primary and metastatic melanoma compared to benign nevi. This could be due to the enhanced activity of sialyltransferases, including ST3GAL1, which has been associated with melanoma migration and invasion ^12^. Only a handful of studies have examined ST3GAL1 function in human cancer. ST3GAL1 is overexpressed in ovarian ^54^ and breast cancers ^55^, and associated with poor prognosis in renal cancer ^56^ and glioblastoma ^57^. Induced expression of ST3GAL1 in mouse mammary cells promotes tumorigenesis ^58^. ST3GAL2 is even less studied in cancer, although recently, overexpression of both the enzyme and a corresponding sialylated glycolipid SSEA-4 were associated with chemotherapy-resistant breast and ovarian cancers, correlating with poor outcomes ^59^.

Our two parallel approaches (i.e., genetic screens and lectin profiling) provided multiple interesting candidates with a possible role in melanoma growth. We applied an additional filter based on transcriptomics data analysis to identify the most robust candidate glycogenes potentially modulating melanoma cell survival. Altogether, our transcriptomics, lectin fluorescence, and immunohistochemistry validation analysis in independent cohorts support a role for ST3GAL1 and ST3GAL2 generated α-2,3-sialosides in melanoma survival and growth. *ST3GAL1* is partially essential in various cancer cell types with a median essentiality score of −0.2 as observed by DepMaP CRISPR perturbation data. While *ST3GAL1* and *ST3GAL2* are essential genes for melanoma cells, their silencing does not abrogate the survival of normal melanocytes such as NHEM-A or cell lines from other cancer types (Fig. S5D). These results support the value of an integrated approach to identifying specific glycogenes and corresponding glycan changes contributing to melanoma transformation. Moreover, our data suggest that ST3GAL1 and ST3GAL2 upregulation in melanoma is either directly pro-oncogenic or an event positively selected for during tumorigenesis to support survival or growth during neoplastic transformation.

Glycans can influence cancer cells properties by altering the glycosylation of adhesion molecules, membrane receptors, etc. To dissect the role of α-2,3 sialylation on growth-related programs of melanoma, we identified sialylated proteins and the mechanisms by which α-2,3 sialylation impacts them. MAA lectin-based glycoproteomic analysis of melanoma cell lines identified a set of α-2,3 sialylated membrane proteins, enriched in factors involved in cell growth, among other biological processes. These include TFR1 or transferrin receptor, a membrane glycoprotein which can import iron by binding a plasma glycoprotein, transferrin (TF) ^60^, and involved in tumor progression and abundantly expressed in liver, breast, lung, and colon cancer cells ^61–64^; ApoE which suppresses melanoma progression and metastasis ^65, 66^; SLC1A4, a glutamine transporter which promotes apoptosis in Hepatocellular Carcinoma^67^ and decreases growth of prostate cancer cells ^68^; SERPINH1, a serine proteinase inhibitor which contributes to gastric cancer progression ^69^; and LGALS3BP, a galectin binding protein which is associated with poor prognosis in several human cancers ^70, 71^. Some of these proteins have been previously shown to be glycosylated ^72^. We focused our studies on SLC3A2 (CD98 heavy chain), a type II transmembrane protein that covalently links to one of several L-type amino acids (AA) transporters (light chains) to form large, functionally heterodimeric neutral amino acid transport systems ^73, 74^. CD98hc plays a central role in AA and glucose cellular nutrition, redox homeostasis, and nucleotide availability, all key for cell proliferation ^75^. In addition to playing a role in amino acid transport, CD98hc has been shown to associate with β_1_ and β_3_ integrins and regulate integrin signaling, which is in turn involved in cell adhesion, fusion, proliferation, and growth^76, 77^. Multiple reports suggest that overexpression of CD98hc stimulates FAK and AKT phosphorylation ^77–79^, and deletion of CD98hc impairs integrin signaling ^41^.

We found that CD98hc stability is dependent upon ST3GAL1/2-mediated sialylation. Indeed, de-sialylated CD98hc degraded much faster than sialylated CD98hc. Moreover, CD98hc silencing reduced melanoma cell proliferation, and CD98hc ectopic overexpression rescued the antiproliferative effect of ST3GAL1 or ST3GAL2 depletion (Fig. 5D-E). As CD98hc interacts with integrins and activates PI3K/AKT signaling, our data suggest that α-2,3 sialylation by ST3GAL1 and ST3GAL2 contributes to survival by enhancing CD98hc stability in melanoma cells. A possibility is that CD98hc sialylation influences protein export and membrane localization and, indirectly, protein stability. The rapid degradation of de-sialylated CD98hc may be a consequence of improper folding, inefficient binding to a chaperon, or changes in post-translational modifications resulting in protein cleavage or degradation. In fact, LAT1/CD98hc has been found among the multiple transmembrane protein clients of the CRBN-HSP90 co-chaperon complex ^80^.

Overall, our work reveals *ST3GAL1* and *ST3GAL2* as robust pro-survival genes in melanoma, highlighting an essential role of α-2,3 sialylated glycans in melanoma biology and pathogenesis. Moreover, our data support the function of α-2,3 sialylated glycans in the modulation of the activity of CD98hc, an α-2,3 sialylated transmembrane glycoprotein required for melanoma survival (Fig. 6). Our findings may have therapeutic applications in melanoma by targeting melanoma-specific sialylated glycopeptide/s of CD98hc.

## Methods

### Cell Lines and Cell Culture

131/4-5B1 (hereafter 5B1) melanoma cells, A549 non-small cell lung cancer (NSCLC) cell line, and HEK293T cells (for lentivirus production) and were grown in DMEM with 10% FBS, 1 mmol/L sodium pyruvate, 4 mmol/L L-glutamine, 25 mmol/L D-glucose, 100 units/mL penicillin, and 100 μg/mL streptomycin. MeWo was cultured in EMEM with 10% FBS, 1 mmol/L sodium pyruvate, 4 mmol/L L-glutamine, and 25 mmol/L D-glucose. Melanocytes NHEM was purchased from Promocell, cultured in Melanocyte growth media (Promocell), and dissociated for harvest using Detach Kit (Promocell). Cell lines were maintained in a 5% CO_2_ incubator at 37°C and were routinely tested for Mycoplasma contamination. Low-passage melanoma STCs, derived in the Osman laboratory as describe^32^, were grown in Dulbecco’s Modified Eagle Medium (DMEM) with 10% FBS, 1 mmol/L sodium pyruvate, 4 mmol/L L-glutamine, 25 mmol/L D-glucose, 1% nonessential amino acids (NEAA).

### Clinical Specimens

Human melanoma specimens (nevi, primary, metastatic) were collected at the time of surgery (Table S2). Approval to collect specimens was granted by New York University Institutional Review Board protocol number i10362, ‘‘Development of an NYU Interdisciplinary Melanoma Cooperative Group: A clinicopathological database.’’ Informed consent was obtained from all subjects included.

### Mice

NOD/SCID/IL2γR^−/−^ female mice (Jackson labs, Cat# 05557) were used for *in vivo* studies. Experiments were conducted following protocols approved by the NYU Institutional Animal Care Use Committee (IACUC) (protocol number S16-00051).

### Generation of custom shRNA library of glycosyltransferases and *in vivo* shRNA mediated growth screen

To identify glycosyltransferases whose silencing can confer growth inhibition in melanoma, we built a custom shRNA library targeting 199 glycosyltransferases (5-6 shRNAs, a total of 1169 shRNAs) using miR30-adapted sequences (5-6 shRNAs per target) by PCR-cloning a pool of oligonucleotides synthesized on 55k customized arrays (Agilent Technologies) using a well-established system ^84–87^. The library was sub-cloned in TRMPV-Neo using 12 independent sub-pools, each consisting of 100 shRNAs, to ensure that shRNA representation was not lost after grafting the tumor cells *in vivo*. Sequence verification was performed by random screening and Sanger sequencing of individual clones from each subpooled library, as well as Illumina sequencing of the final library. We checked that library purity (percentage of non-mutated shRNAs) is high, around 90%. We found that all the shRNAs are represented in the corresponding libraries, and no cross-contamination was found.

### *In Vivo* shRNA Mediated Screen and HiSeq

Each pool of the library was transduced into MeWo melanoma cells stably expressing rtTA (Addgene #26429) using conditions that predominantly lead to a single retroviral integration and represent each shRNA in a calculated number of 9,600 cells (a total of 1.2 million cells at infection, 8% transduction efficiency, in technical triplicates). For viral library production, GP2 cells were plated on 100 mm tissue culture plates precoated with collagen at about 50% confluency 24 hours before transfection. Transfection was performed with a total of 12 μg of library and 8 μg of VSV-G using Lipofectamine 2000 (Invitrogen) overnight, and media was exchanged for DMEM with 10% Tet-free FBS (Clontech). Supernatants were collected and 0.45 μm filtered at 48 and 72 hours post-transfection and immediately used for infection of MeWo-rtTA target cells in the presence of 8 μg/ml of polybrene (Fisher). Transduced cells were selected for ten days using 600 μg/ml G418 (Invitrogen); at each passage, more than ten million cells were maintained to preserve library representation throughout the experiment. Before induction, time point zero (T0) samples were obtained (one million cells per replicate), and cells were subsequently cultured with 600 μg/ml G418 and 2 μg/ml doxycycline to induce shRNA expression. We combined two sub-pools of transduced MeWo cells in equal numbers before injecting them subcutaneously. Briefly, cells were resuspended in sterile PBS at a concentration of 2 x 10^6^ cells per 150 ml, aliquoted into Eppendorf tubes (150 ml) and maintained on ice until injection. Immediately before injection, cell aliquots were mixed with 150 ml Matrigel (Becton Dickinson). Cell/Matrigel (1:1) suspensions were injected subcutaneously in the right flank of NOD/Shi-scid/IL-2Rgamma null (NSG) 8-week-old female mice (n=10 for each group). Each sub-pool of cells was injected in mice in triplicates. Primary tumors were resected five weeks post-injection. Genomic DNA from 5 weeks post-injection tumors was isolated by two rounds of phenol extraction using PhaseLock tubes (5 prime) followed by isopropanol precipitation. Deep-sequencing template libraries were generated by PCR amplification of shRNA guide strands ^86^. Libraries were analyzed on an Illumina HiSeq at a final concentration of 10 pM; 18 nucleotides of the guide strand were sequenced using a custom primer (miR30 EcoRISeq, TAGCCCCTTGAATTCCGAGGCAGTAGGCA). Sequence processing was performed using a customized Galaxy platform ^88^. For each shRNA and timepoint, the number of matching reads was normalized to the total number of library-specific reads per lane. For scoring of enrichment/depletion of each shRNA, an average value for each timepoint divided by the respective input was averaged, and log10 fold change was calculated. We applied various cutoffs in a sequential manner to identify shRNAs depleted or enriched, as shown in fig S1A.

### Lectin Microarray Printing, Hybridization and Analysis

Lectins were purchased from E.Y. Laboratories or Vector Laboratories with the following exceptions: recombinant CVN, SVN and Griffithsin were gifts from B. O’Keefe (NCI, Frederick, MD); TJA-I and TJA-II were from NorthStar Bioproducts. Printing, hybridization, and data analysis were performed as previously described ^89^. Our data were normally distributed as determined by the Lilliefors test in MATLAB. Lectins were excluded from the analysis if they did not meet our minimum threshold for activity.

### FFPE Nevi, Primary, and Metastatic Melanoma Sample Deparaffinization, Protein Extraction and Labeling

Formalin-fixed paraffin-embedded (FFPE) tissues from benign nevi (n= 10) and patients with primary melanoma (n = 18), LN metastasis (n = 22), subcutaneous metastasis (n = 21) brain metastasis (n = 18) were analyzed on our lectin microarrays. In brief, melanoma samples were fixed in 10% neutral buffered formalin. To analyze tumor samples, hematoxylin and eosin-stained slides were reviewed to ensure enough viable tumors. Unstained cut sections mounted on slides were macro dissected to remove contaminating normal cells, and two 20 µm sections of each FFPE tissue were scraped into a 1.5-ml microcentrifuge tube. One mL xylene and 200 ml ethanol were added to a microcentrifuge tube, incubated for 15 min at room temperature, centrifuged at 10,000 x g for 2 min, and the supernatant was removed. The deparaffinized tissue pellets were then rehydrated with a graded series of ethanol. One mL of 95% ethanol was added to each tissue pellet, incubated for 10 min at room temperature, and centrifuged at 10,000 x g for 2 min. This was followed by rehydration with 70% ethanol, spun down to remove excess supernatant, and allowed samples to dry at room temperature. The rehydrated tumor tissue and cell sections were re-suspended in 200 ml 10 mM sodium citrate buffer (pH 6.0) and incubated at 95° C for 60 min. The supernatant was removed, and the tissue was washed with PBS. Pellet was solubilized with 250 ml (100 ml for smaller pellets) Cy buffer (0.1 M NaHCO_3_, pH=9.3) containing 0.5% NP40 (Nonidet P-40). The sample was sonicated gently (70% power, total time 1 min [10 s on, 10 s off]) and incubated on ice for 60 min. The sample was centrifuged at 12,000 x g for 5 min at 4° C. Supernatant was collected for analysis. Total protein concentration was quantified by BCA assay. Samples were labeled with NHS-Cy3 or -Cy5 and analyzed on our lectin microarrays in dual color using standard protocols^89^. The reference sample was a mixture of 3 nevi, 6 primary, and 6 metastatic samples. Normalized data were subjected to hierarchical clustering using the Pearson correlation coefficient (R) as the distance metric and average linkage analysis to generate heat maps.

### RNA Extraction and Real-Time qPCR

Total RNA was extracted from samples (miRNeasy mini kit; Qiagen), quantified using Nanodrop 8000, and stored at −80° C. 500 ng of RNA were reverse transcribed using TaqMan RT reagents (Applied Biosystems) with random hexamers following manufacturer’s recommendations. Transcripts were quantified by real-time qPCR using Power SYBR Green PCR MasterMix (Applied Biosystems) in StepOne System (ABI). Primers were designed by using PrimerSelect. Some qPCR primers were purchased from Origene (**Table S3**). Cycle threshold values were normalized to those of the housekeeping genes GAPDH. The average for three biological replicates was plotted as relative transcript abundance.

### Plasmids

pLKO.1 plasmids targeting human ST3GAL1, ST3GAL2, CD98hc and a non-targeting control were purchased from Sigma. DOX inducible PLKO plasmids were subcloned using shRNA and scramble sequences, as shown in table S3. CD98hc overexpression plasmid pCMV6-Myc-DDK was purchased from Origene.

### Viral Production

4×10^6^ HEK293T cells were seeded per 10 cm tissue culture dish and incubated overnight at 37°C and 5% CO2 for 16-20 hr. After seeding, HEK293T was co-transfected with lentiviral expression constructs (12 µg), viral packaging plasmid (psPAX2, 8 µg), and viral envelope plasmid (pMD2.G, 4 µg) using Lipofectamine 2000 (Invitrogen) following manufacturer’s recommendations. Viral supernatant was collected and 0.45 µm filtered at 72 hr post-transfection and stored at 4° C for short-term use (1-5 days) or −20 C for long-term storage (5-30 days).

### Viral Transduction

Target cells were seeded and incubated overnight prior to infection. The medium was replaced with 1:2 diluted viral supernatant supplemented with eight µg/mL polybrene and incubated for 6-8 hr, followed by replacement with growth medium. Control and shRNA cells were selected using puromycin (2 µg/mL) or neomycin (500 µg/mL) prior to use in experiments.

### *In Vitro* Proliferation Assay

Cells were seeded at 5000 cells/well in 96-well plates. Cell proliferation was measured using CellTiter-Glo 2.0 Luminescent Cell Viability Assay (Promega) following the manufacturer’s instructions.

### Caspase activity assay

Cells were treated with or without a pan-caspase inhibitor (10 nM, QVD) 24 hr post-lentiviral infection. Cells were then rinsed 1x with PBS and resuspended in 1X lysis buffer (20 mM HEPES/NaOH, pH 7.2, 10% sucrose, 150 mM NaCl, ten mM DTT, five mM EDTA, 1% Igepal CA-630, 0.1% CHAPS, and 1X EDTA-free Complete protease inhibitor mixture (Roche)). Lysates were cleared by centrifugation at 16,000x*g* for 5 min, and supernatants were quantified by the Lowry method (Bio-Rad). Assays were performed in triplicate using 25 μg of protein in lysis buffer supplemented with 25 μM of the fluorogenic substrate Ac-DEVD-afc. Plates were read in a fluorimeter using a 400 nm excitation filter and a 505 nm emission filter.

### Immunohistochemistry (IHC)

Unconjugated anti-human ST3GAL1 and ST3GAL2, raised against human ST3GAL1 (Sigma HPA040466) and ST3GAL2 (Abcam ab96028), were used for IHC. Antibody optimization was performed on formalin-fixed, paraffin-embedded 4-micron composite tissue microarrays (TissueArray.Com) containing normal and tumor: skin, stomach, prostate, liver, and pancreas. IHC parameters were validated on formalin-fixed, paraffin-embedded 4-micron human hippocampus with cells clearly expressing cytoplasmic-membrane staining considered positive. Chromogenic IHC was performed on a Ventana Medical Systems Discovery XT instrument with online deparaffinization using Ventana’s reagents and detection kits unless otherwise noted (Ventana Medical Systems Tucson, AZ USA). AMIGO2 was antigen-retrieved in Ventana Cell Conditioner 1 (Tris-Borate-EDTA) for 20 min. Antibody was diluted 1:50 in PBS and incubated for 6 hr at room temperature. Primary antibody was detected with goat anti-mouse, horseradish peroxidase-conjugated multimer incubated for 8 min. The complex was visualized with alpha-naphthol pyronin incubated for 8 min. Slides were washed in distilled water, counterstained with Hematoxylin, dehydrated, and mounted with permanent media. Negative controls were incubated with diluent instead of primary antibody. The staining score was performed in a blinded manner. The IHC score was calculated using staining intensity (0, 1, and 2) x percentage of positive cells.

### Lectin Fluorescence Microscopy

Lectin fluorescence was performed on 4 mm formalin-fixed, paraffin-embedded tissue sections using diCBM40-Cy5 lectin. Both lectins and secondary antibodies were incubated for 1 hr each. DAPI staining was used to determine the nuclei morphology. Fluorescent slides were stored at −20° C until imaging. Stained slides were imaged by Hamamatsu fluorescent slide scanner, and images were extracted using NDPIS software. Lectin fluorescence microscopy for cultured cells was performed as previously described ^90^.

### Western Blotting

Cells were lysed in cold RIPA buffer supplemented with protease inhibitors. Equal amounts of protein were resolved by 4-12% Bis-Tris/ PAGE, transferred onto PVDF membranes, and blocked in buffer [5% (wt/vol) milk, TBST (TBS, pH 7.4, 0.05% Tween-20), 1 hr, room temperature]. Primary antibodies were diluted in blocking buffer, as follows: ST3GAL1 (1:1000, Sigma HPA040466), ST3GAL2 (1:1000, ab96028), CD98hc (1:5000, Abcam ab244356), TFR1 1:5000, Abcam ab269513), PAPR (1:4000, Cell signaling, 9542), Caspase-3 Antibody (1:4000, Cell signaling, #9662), Cleaved-caspase3 (1:2000, Cell signaling, #9661), (GAPDH (1:5,000; Abcam, ab8245); α-Tubulin (1:5000; Sigma, T9026). Secondary antibodies were a-mouse or α-rabbit-HRP (1:5,000; Bio-Rad). Blots were developed using Clarity Western ECL Blotting Substrate (BioRad) and imaged in the LICOR Odyssey Fc imaging system.

### MAA Chromatography

For lectin enrichment assays, cells were lysed with lysis buffer containing 1% Nonidet P-40. 1000 mg of lysate was mixed with 100 µl of biotinylated MAA lectin, and a volume made up to 1000 ml with PBS containing 1mM MnCl_2_, 1mM MgCl_2_, and 1mM CaCl_2_ and incubated with rotation at 4° C overnight. Then, 60 µl of a 1:1 suspension of agarose coupled streptavidin was added, and incubation was continued for 4 hr. The beads were washed five times with PBST (Tween, 0.05%) buffer and subsequently extracted with SDS-PAGE sample buffer at 95° C for 5 min. The samples were separated by 4-12% Bis-Tris/PAGE and subjected to immunoblotting with α-CD98hc and α-TFR1 antibodies.

### GO Analysis

GeneOntology Enrichment Pathway analysis was performed using DAVID to determine biological process categories enriched in MAA-enriched alpha 2,3 sialylated glycoproteins. The input gene lists were generated from overlapping α-2,3 sialylated proteins in 3 melanoma cell types consisting of 44 proteins.

### Data mining of human transcriptomics datasets

The log fold changes in gene expression between melanoma or nevi were calculated in the following three Affymetrix transcriptomic datasets: Talantov *et al. ^81^* (GSE3189; 18 nevi and 45 melanomas), Kabbarah *et al. ^82^* (GSE46517; 9 nevi and 104 melanomas) and Scatolini *et al*. ^83^ (GSE12391; 18 nevi and 28 melanomas). Statistical comparisons were made by Student’s *t-testing*. Significant differences in gene expression were considered when *p* < 0.05.

## Supporting information

Supplementary table 3, https://www.origene.com/catalog/vectors/lentiviral-gene-expression-vectors/ps100102/plenti-c-mgfp-p2a-bsd-lentiviral-gene-expre

## Acknowledgments

We thank the NYU School of Medicine Histopathology and IHC Core Laboratories for tissue processing and histology and the NYULMC Proteomics Resource Center for mass spectrometry data. We thank Garrett Cooper (Interdisciplinary Melanoma Cooperative Group, NYU Langone Health) for providing clinical data for melanoma cases. This work was funded by the NIH/NCI R01 CA202027, U54 CA263001. P.A. has been supported by a career development grant Harry J Lloyd charitable trust.

## Supplementary Material

### Supplementary figures

**Fig S1:**
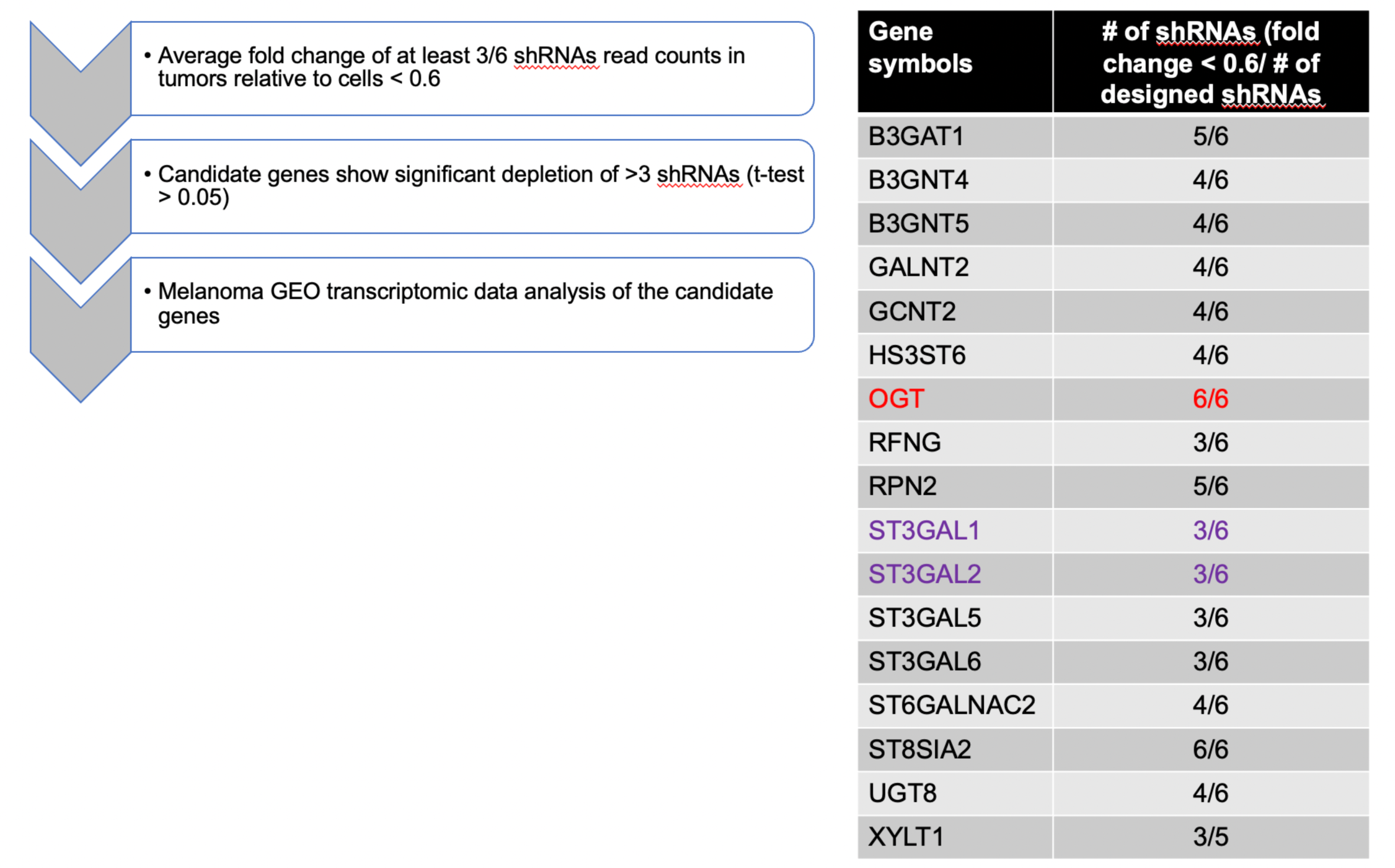
Selection of glycogene candidates using *in vivo* shRNA functional screen (**A**) Stepwise filtering of depleted shRNA. (**B**) List of glycogenes for which # of shRNAs were consistently depleted.

**Fig S2:**
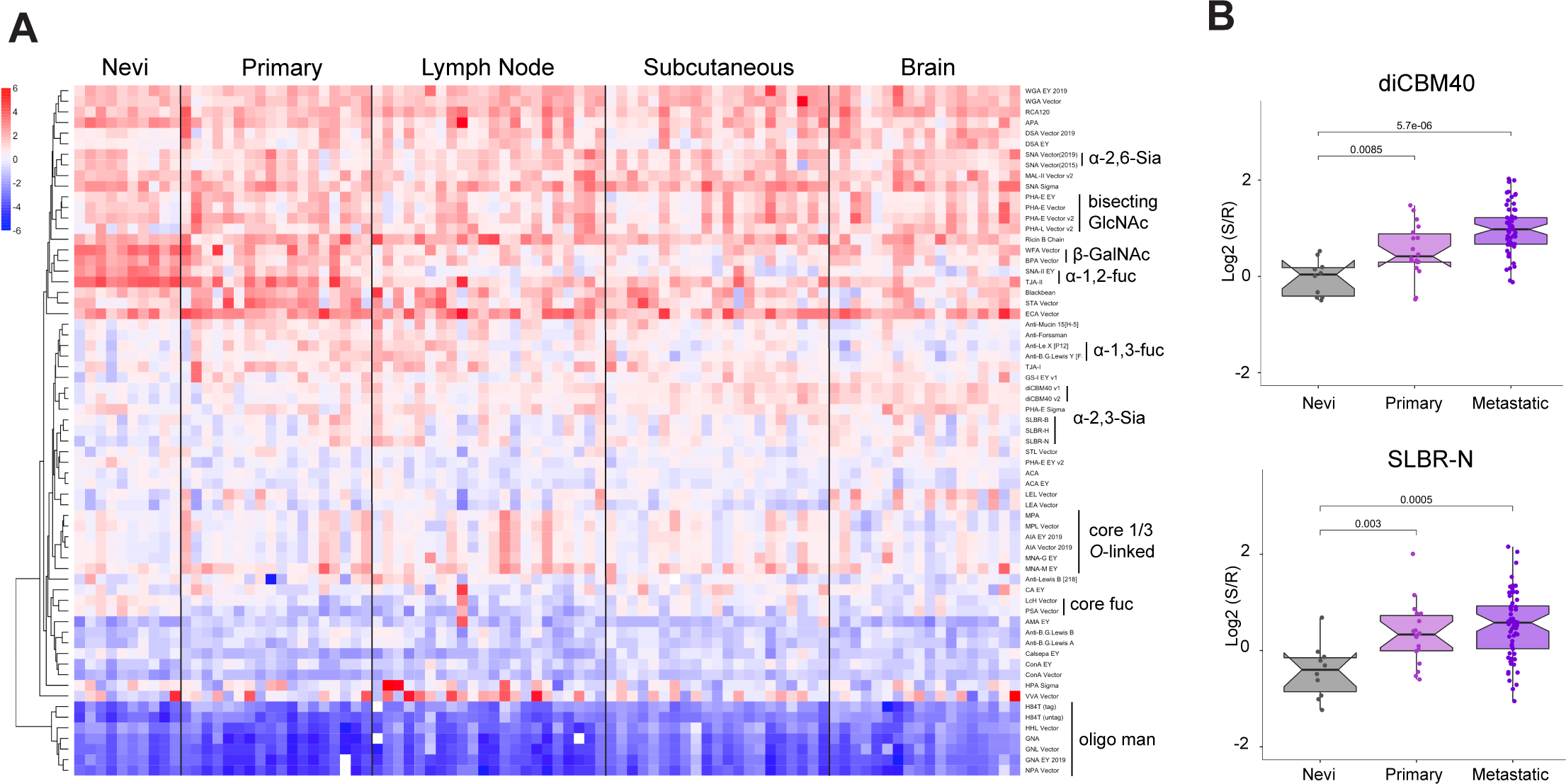
Glycome of melanoma differs from melanocytic nevi. (**A**) Glycomic profile of nevi (n = 10), primary melanoma (n = 18), lymph node metastasis (n = 22), subcutaneous metastasis (n = 21) brain metastasis (n = 18) by ratiometric lectin microarray data. All specimens were micro-dissected from formalin-fixed, paraffin-embedded tissues, and protein levels were determined by the BCA method. Equal amounts (5.0 μg by proteins) of Cy-5 labeled samples, and Cy-3 labeled references were analyzed by lectin microarray. A heatmap of lectin microarray data for all lectins is shown. Median normalized log_2_ ratios (Sample (S)/Reference(R)): red, log_2_(S) > log_2_(R); blue, log_2_(R) > log_2_(S). Lectins were hierarchically clustered using the Pearson correlation coefficient and average linkage analysis. Lectins bindings are highlighted to the right of the heatmap. (**B**) Boxplot analysis of **α**-2,3 sialosides probed by lectins diCBM40 and SLBR-N. Significance was determined using Student’s t-test (two-tailed).

**Fig. S3:**
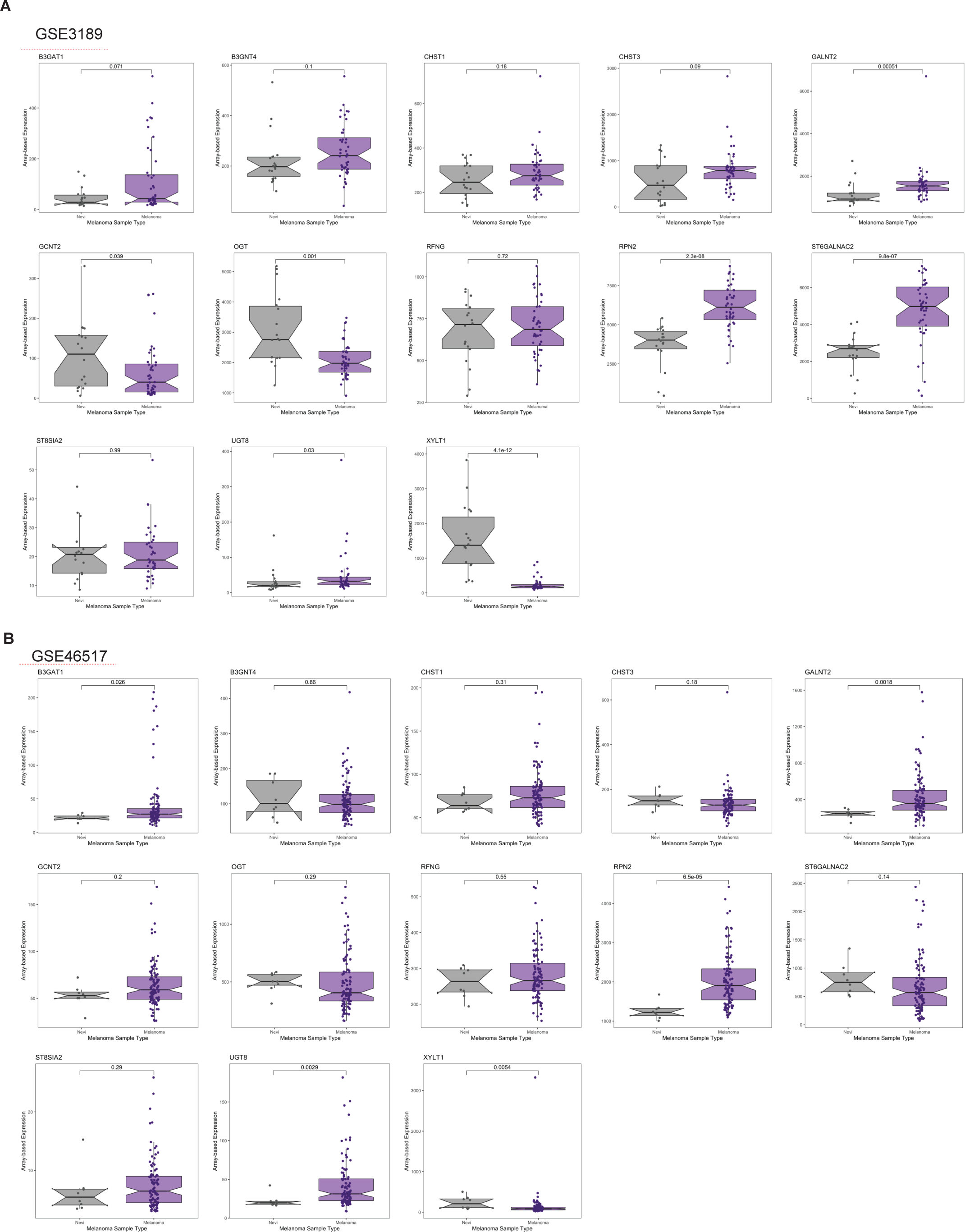
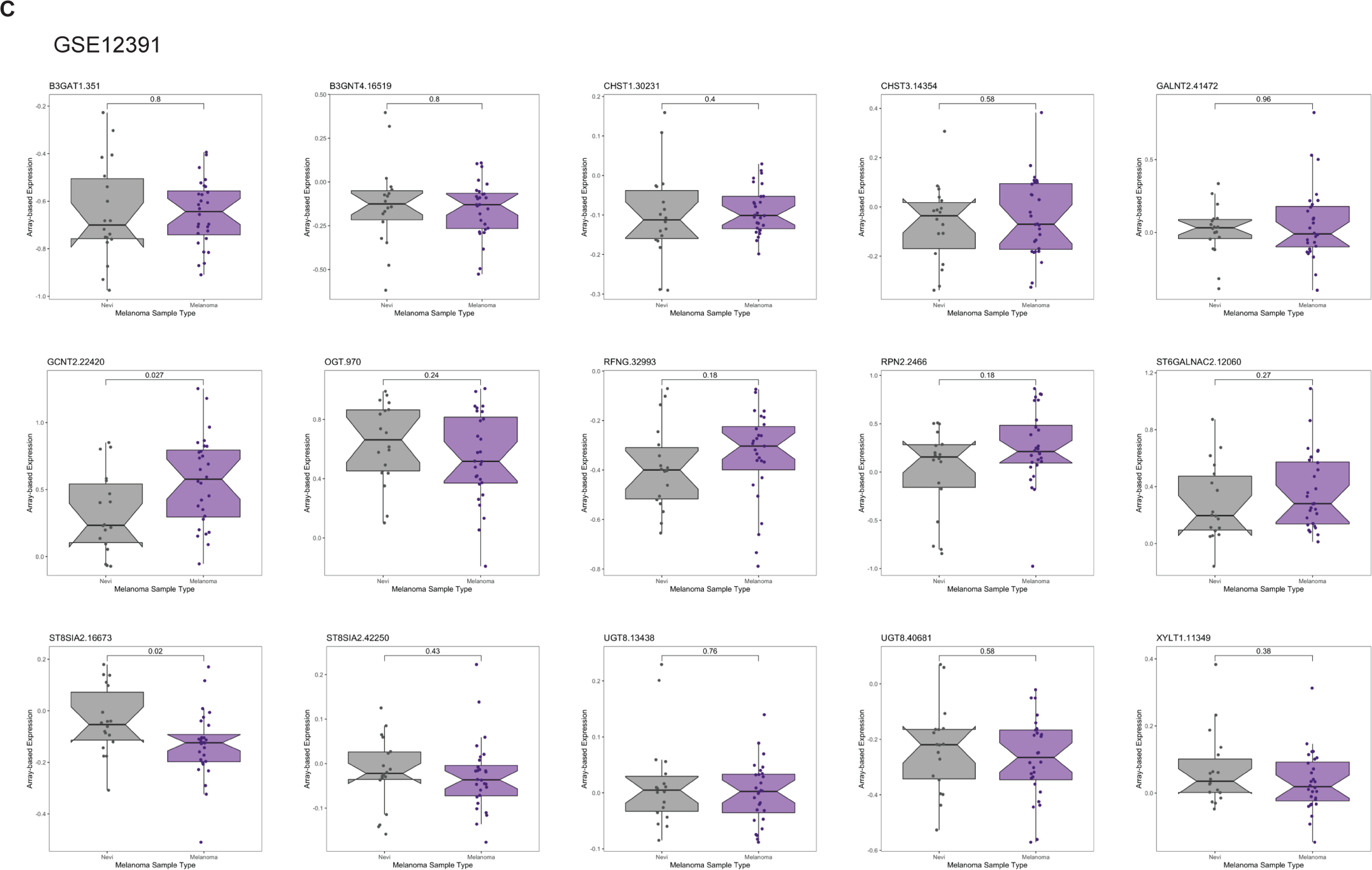
Expression of candidate glycogenes in melanoma relative to nevi. Whisker plot illustrating mRNA expression level in melanoma samples compared to nevi in multiple datasets: **(A)** GSE3189(81), **(B)** GSE46517(82), **(C)** GSE12391 (83). Various glycogene candidates from *in vivo* functional library screening are presented in fig. S1B. Two-tailed unpaired t-test.

**Fig. S4:**
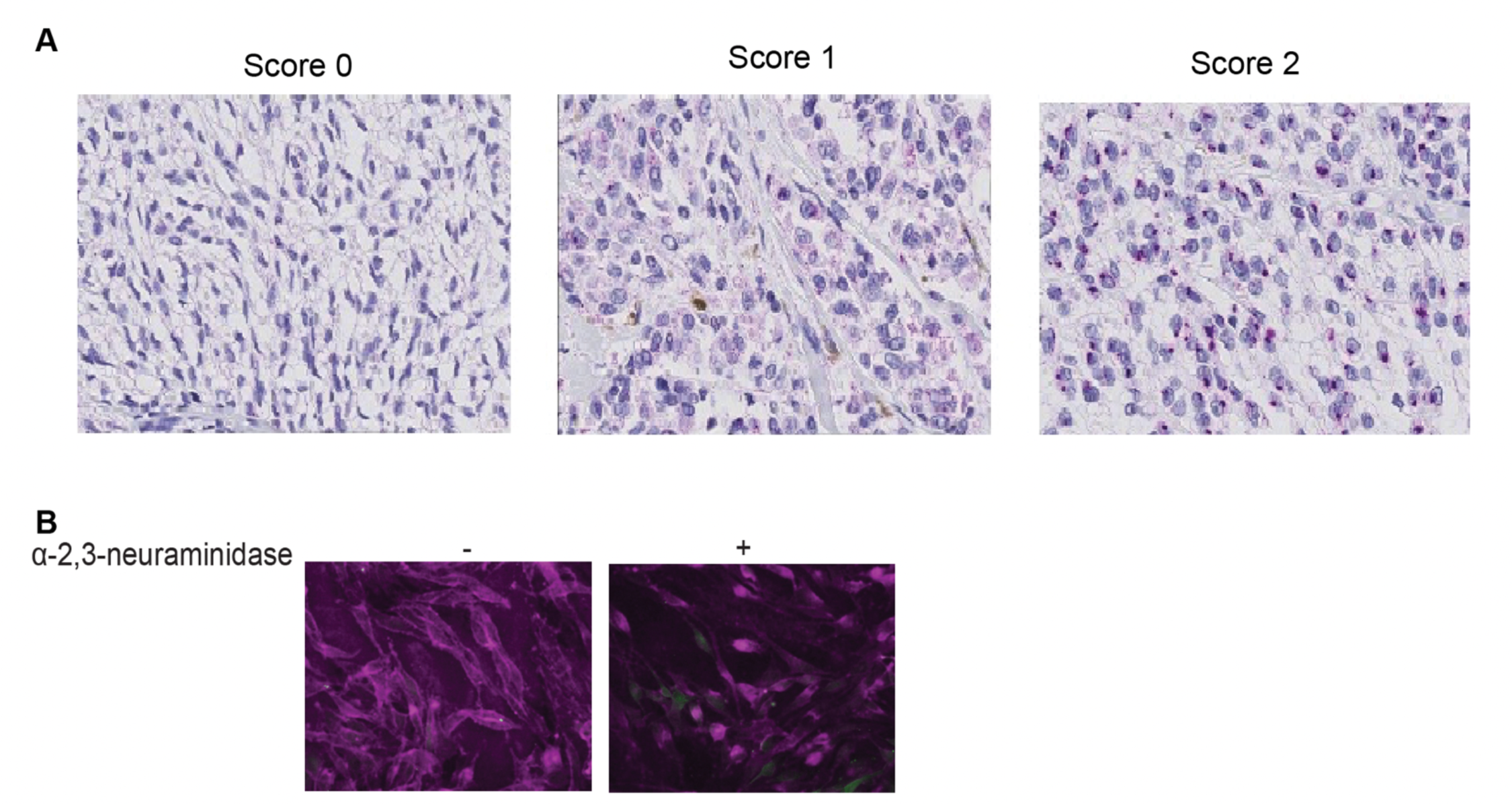
**(A)** Representative ST3GAL1 IHC images depict IHC scoring categories. (**B**) diCBM40 fluorescence microscopy of 5B1 cells with or without α-2,3-neuraminidase (NEB, P0743) treatment. 5B1 cells were incubated with his tagged diCBM40 after neuraminidase treatment and visualized by 6x-His Tag Monoclonal Antibody (HIS.H8), Alexa Fluor™ 647.

**Fig. S5:**
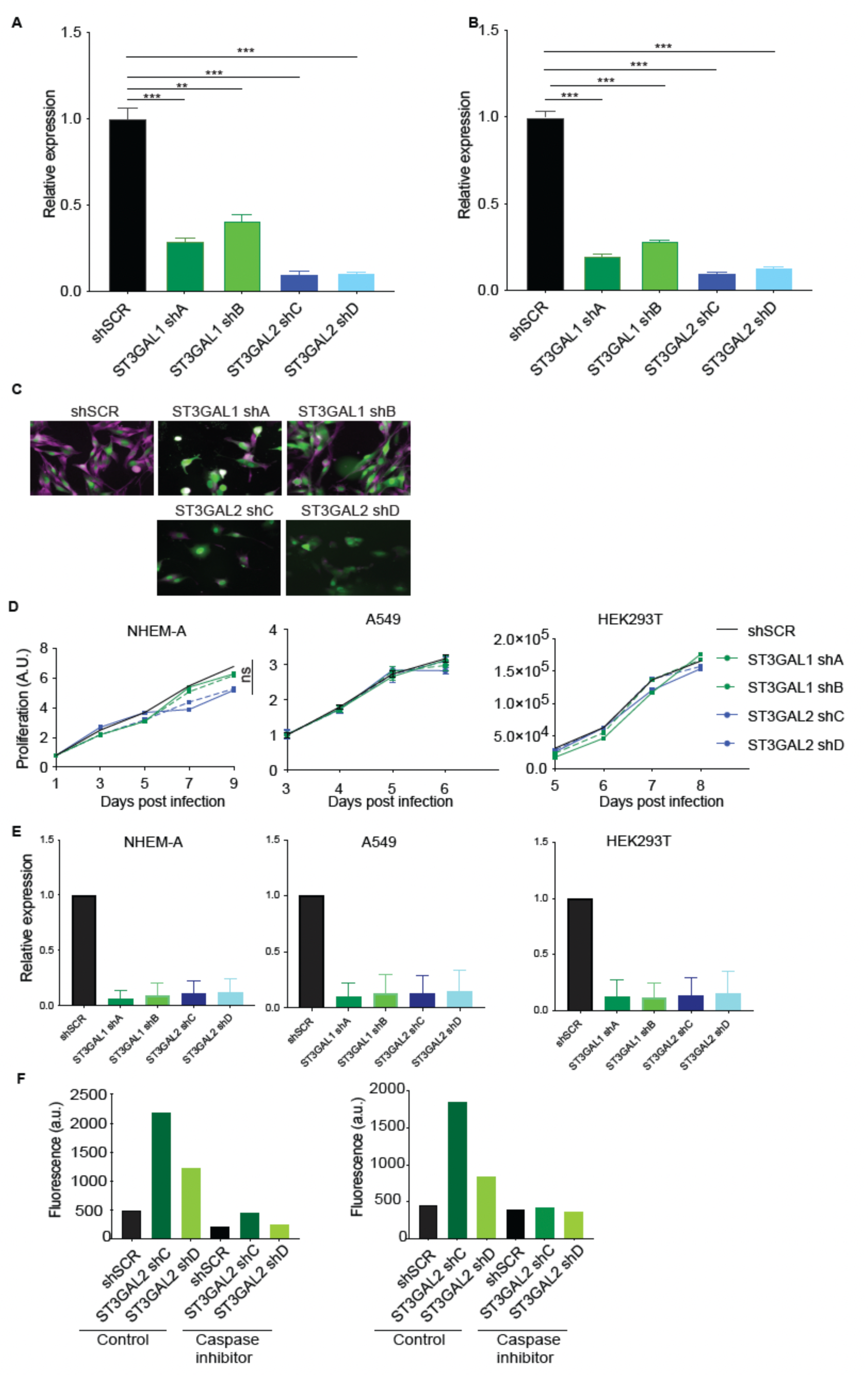
ST3GAL1 and ST3GAL2 are essential for melanoma proliferation *in vitro*. (**A and B**) ST3GAL1 and ST3GAL2 transcript expression in 5B1 and 12-273BM cells stably expressing shRNA targeting ST3GAL1 (shA or shB) or ST3GAL2 (shC or shD) or shSCR were assessed by real-time qPCR. qPCR graph shows average relative expression normalized to *GAPDH*, three replicates per condition, two-tailed unpaired t-tests. qPCR data are representative of three independent experiments. (**C**) diCBM40 fluorescence microscopy of 5B1 cells transduced by ST3GAL1 (shA or shB) or ST3GAL2 (shC or shD) or shSCR. 5B1 cells were visualized by incubating with diCBM40 lectin followed by 6x-His Tag Monoclonal Antibody (HIS.H8), Alexa Fluor™ 647. (**D**) Relative growth curves and (**E**) real-time qPCR of NHEM-A, A549, and HEK293T cells stably transduced with non-targeting scrambled control shRNA (shSCR), *ST3GAL1* shRNAs (shA and shB) and *ST3GAL2* shRNAs (shC and shD). (**F**) FACS analysis of cleaved caspase-3 on melanoma cells, 5B1 (left) and 12-273BM (right) transduced with two independent shST3GAL2 (shC and shD) treated with caspase inhibitor or vehicle.

**Fig. S6:**
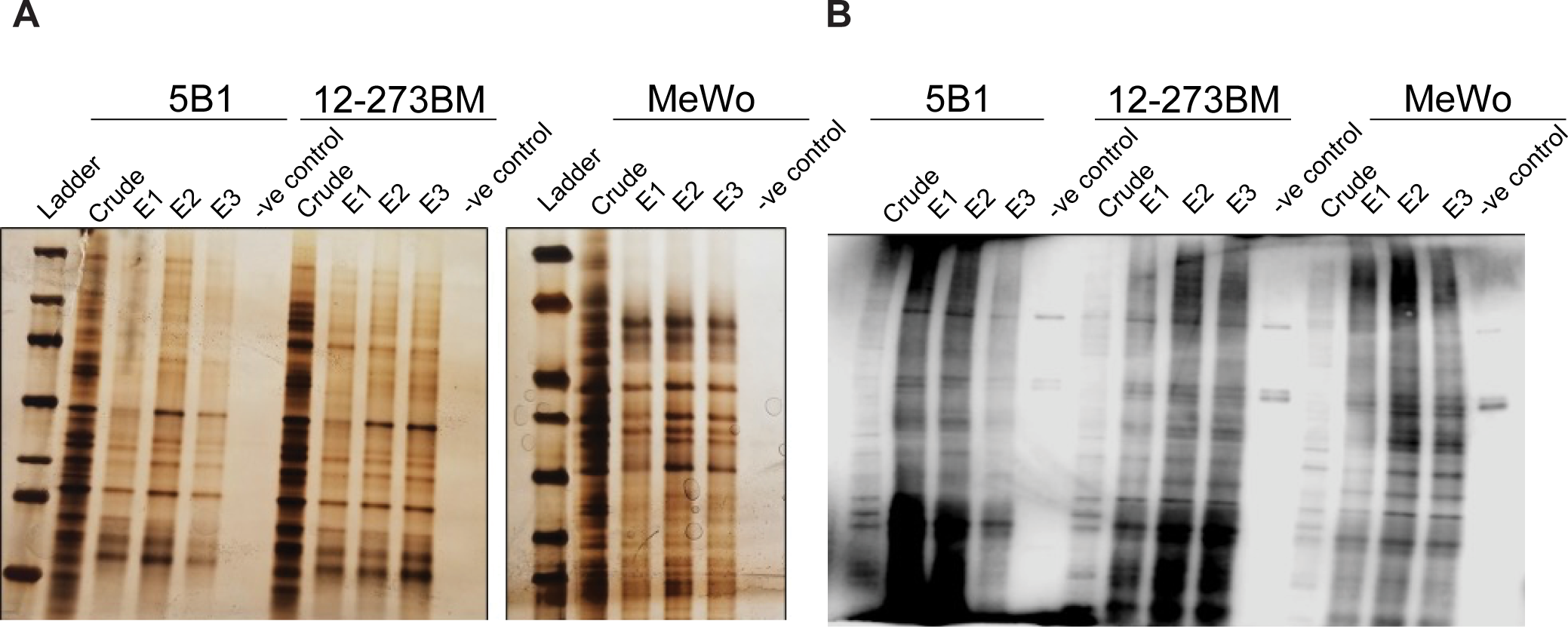
Identification of α-2,3-sialylated glycoproteins in melanoma. Evidence of enrichment of α-2,3 sialylated proteins by **(A)** silver staining and **(B)** MAA lectin blot. An equal volume of MAA-enriched protein was loaded on the gel for silver stain or MAA lectin blot. The left lane shows a molecular weight marker (MW). Samples for each cell line were loaded in 3 biological replicates represented by E1, E2, and E3, along with negative control (without MAA lectin in the enrichment process).

**Fig. S7:**
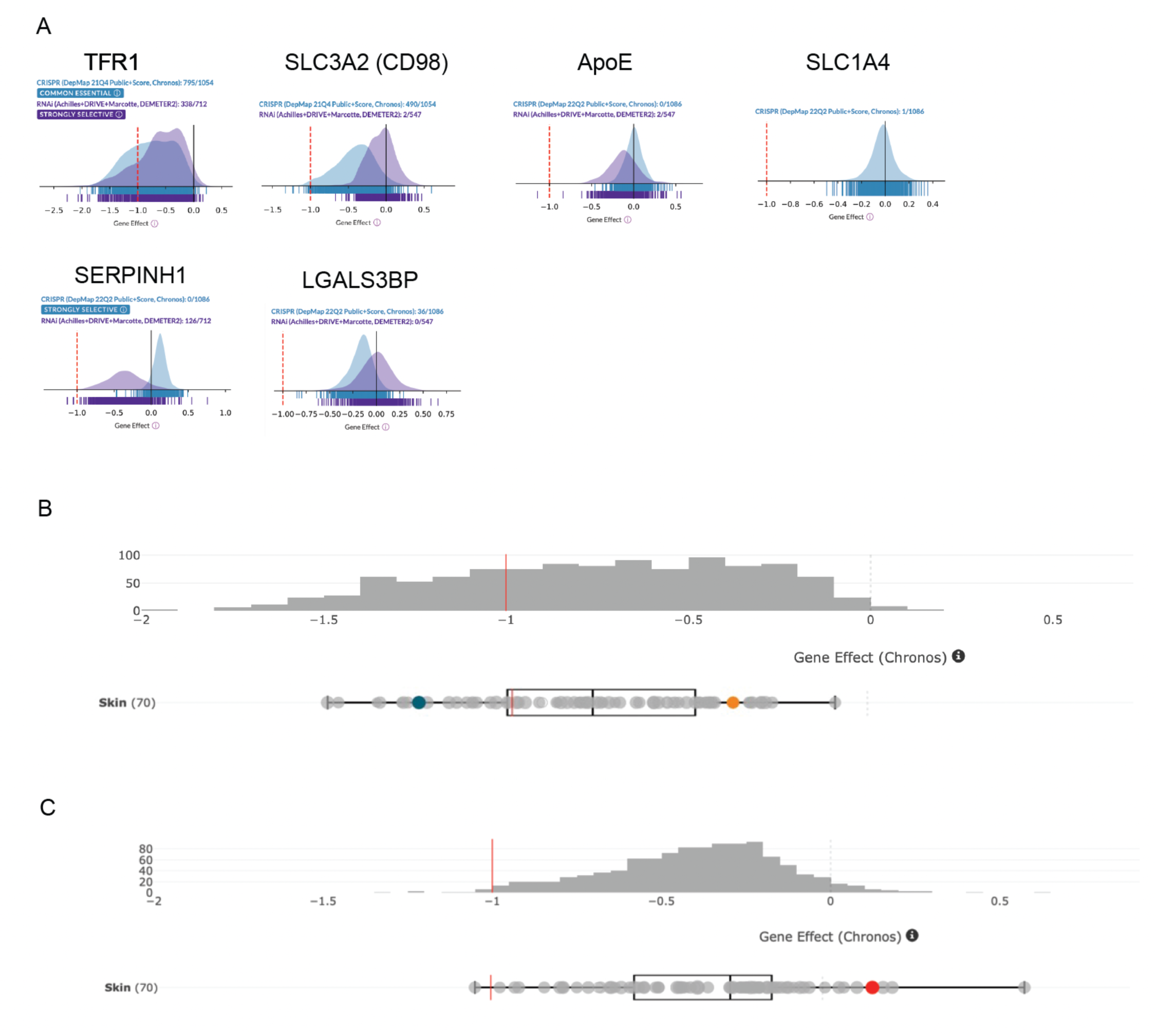
**(A)** Association of sialylated proteins with an essentiality of genes: DepMap data mining of genes for various sialylated glycoproteins in cancer cells. **(B, C)** DepMap data mining of TFR1 and SLC3A2 in melanoma cell lines. Each dot represents a melanoma cell line. Gene effect (Chronos score) is shown. A lower Chronos score means a gene is more likely to be dependent in a given cell line. A score of −1 (red line) corresponds to the median of all common essential genes.

**Fig. S8:**
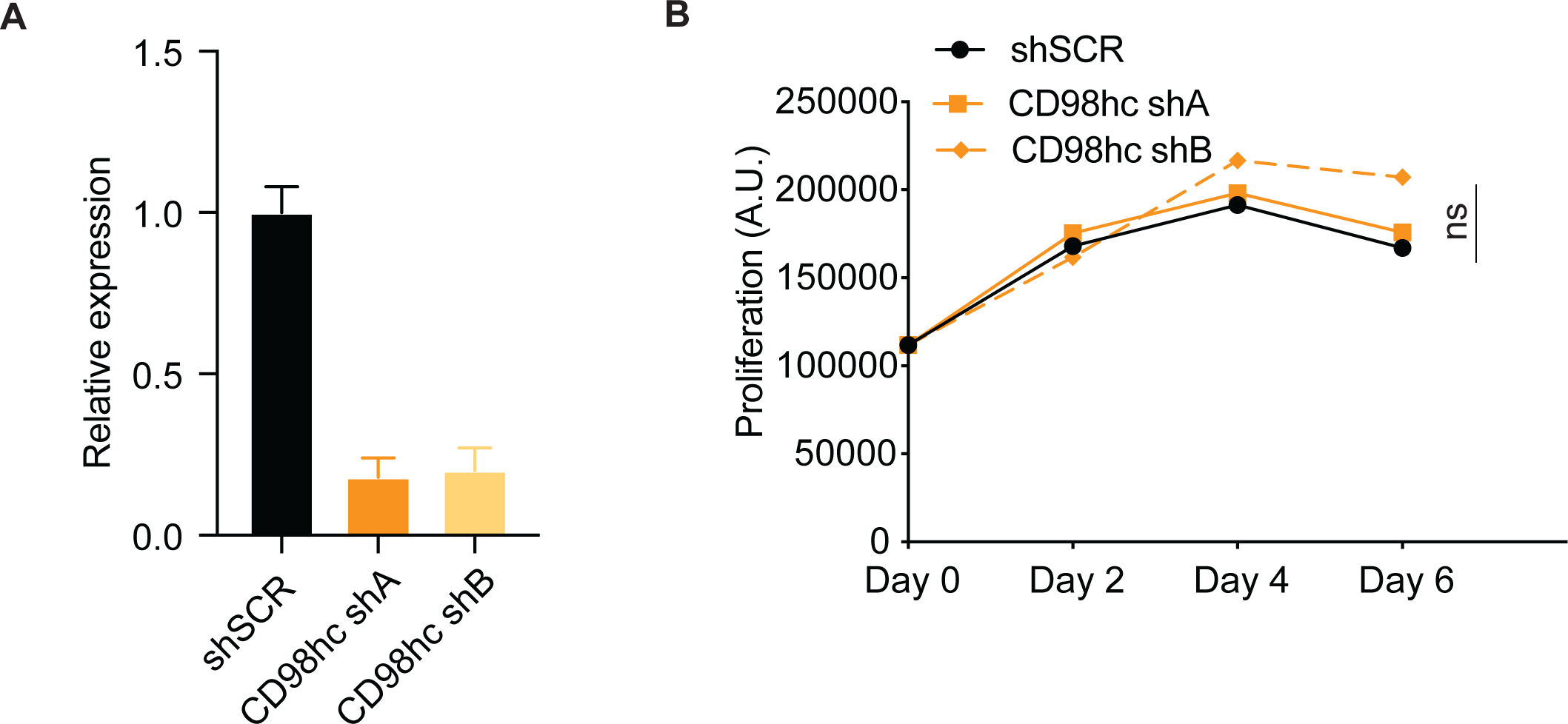
**(A)** SLC3A2 transcript expression in HEK293T cells stably expressing shRNA targeting CD98hc (shA or shB) or shSCR was assessed by real-time qPCR. qPCR graph shows the average relative expression normalized to GAPDH. **(B)** The relative growth curves of HEK293T cells are stably transduced with non-targeting scrambled control shRNA (shSCR) or shRNA targeting CD98hc (shA or shB).

**Fig. S9:**
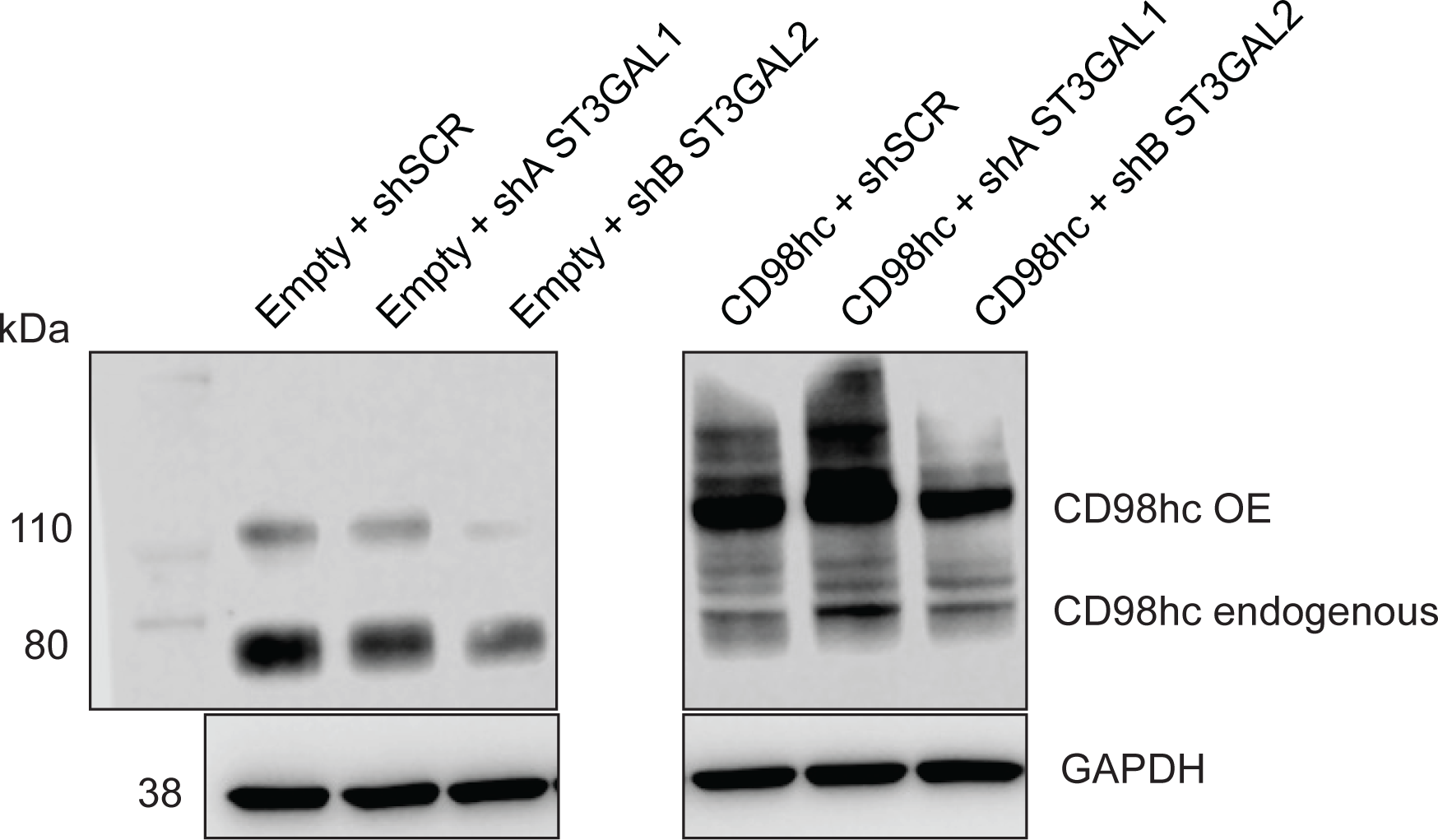
Western blot for CD98hc on lysates from melanoma cells stably overexpressing CD98hc or empty vector and transduced with non-targeting control shSCR or shST3GAL1 or shST3GAL2.

**Fig. S10:**
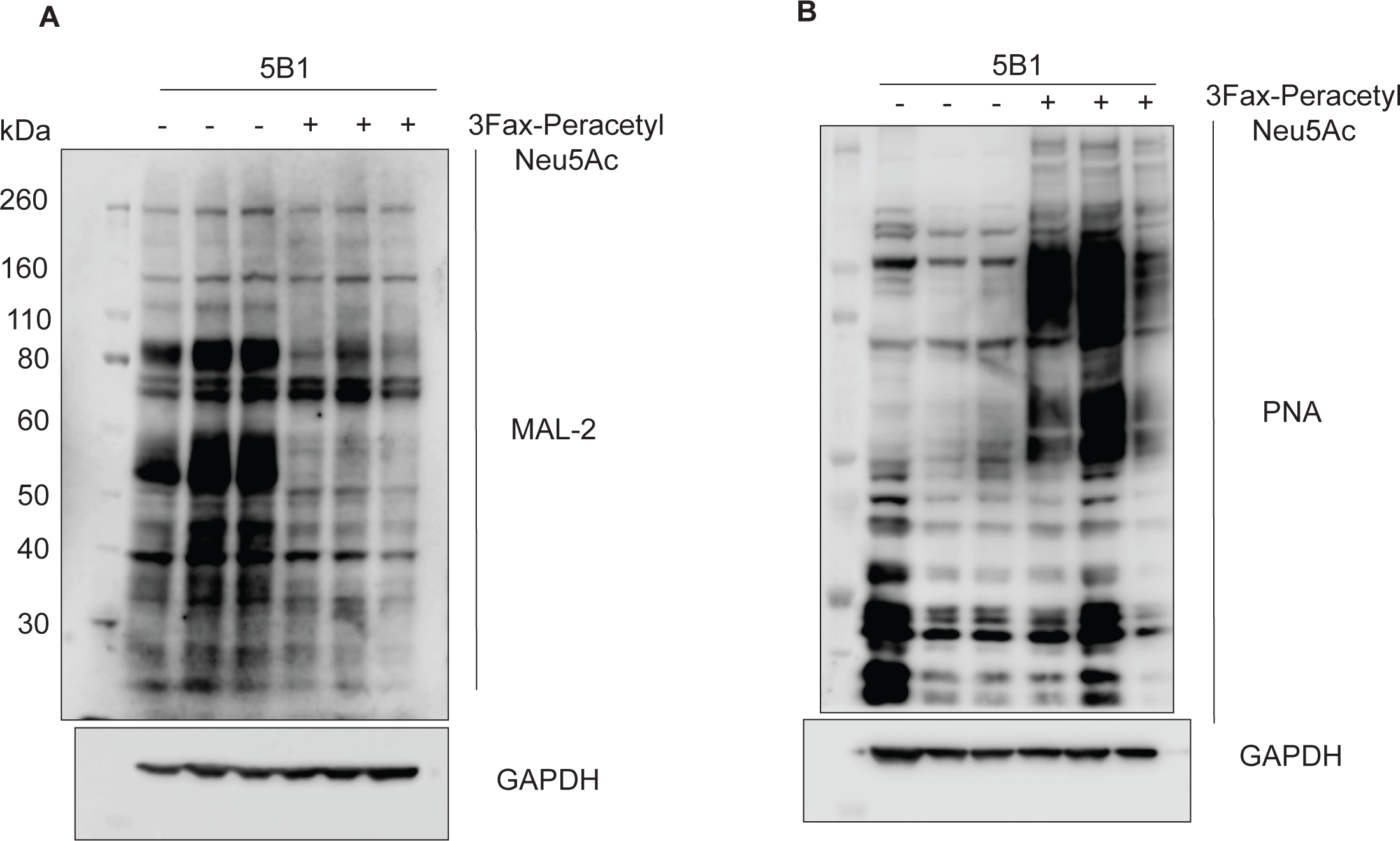
Lectin blot analysis of 5B1 cells treated with sialyltransferase inhibitor 3Fax-peracetylNeu5Ac. **(A)** MAL-2 lectin blot **(B)** PNA lectin blot. GAPDH serves as a loading control.

### Supplementary Tables

**Table S1:**
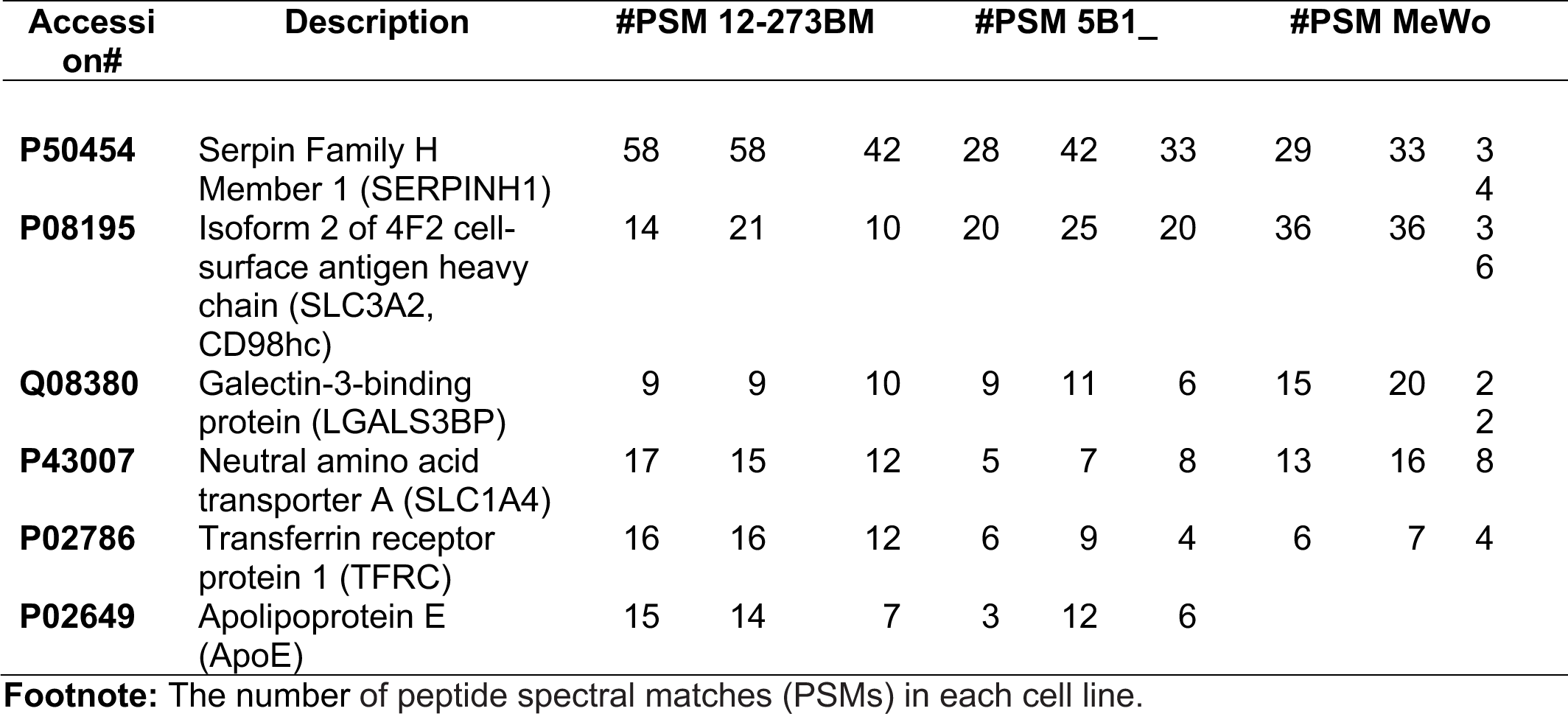
Mass spectrometric analysis of MAA enriched proteins commonly present in the growth category of GO analysis in 5B1, MeWo cells, and 12-273BM STC.

**Table S2:**
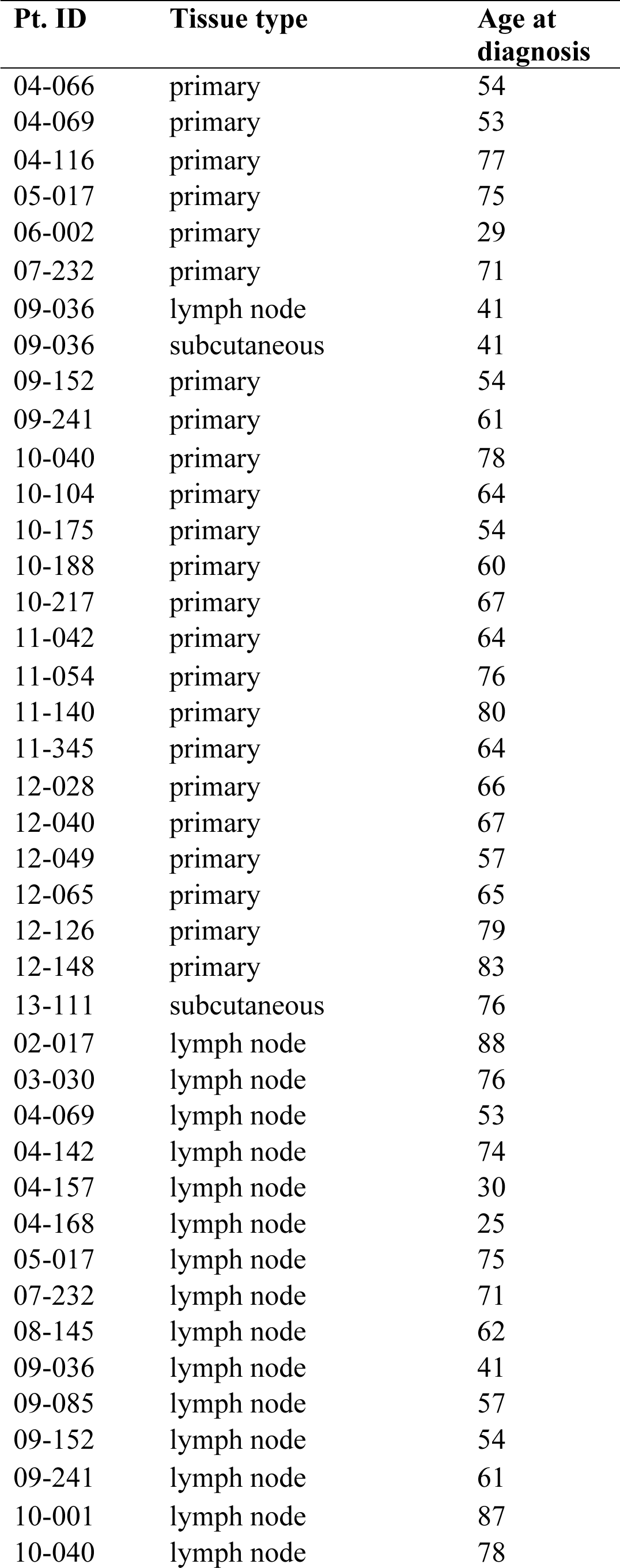
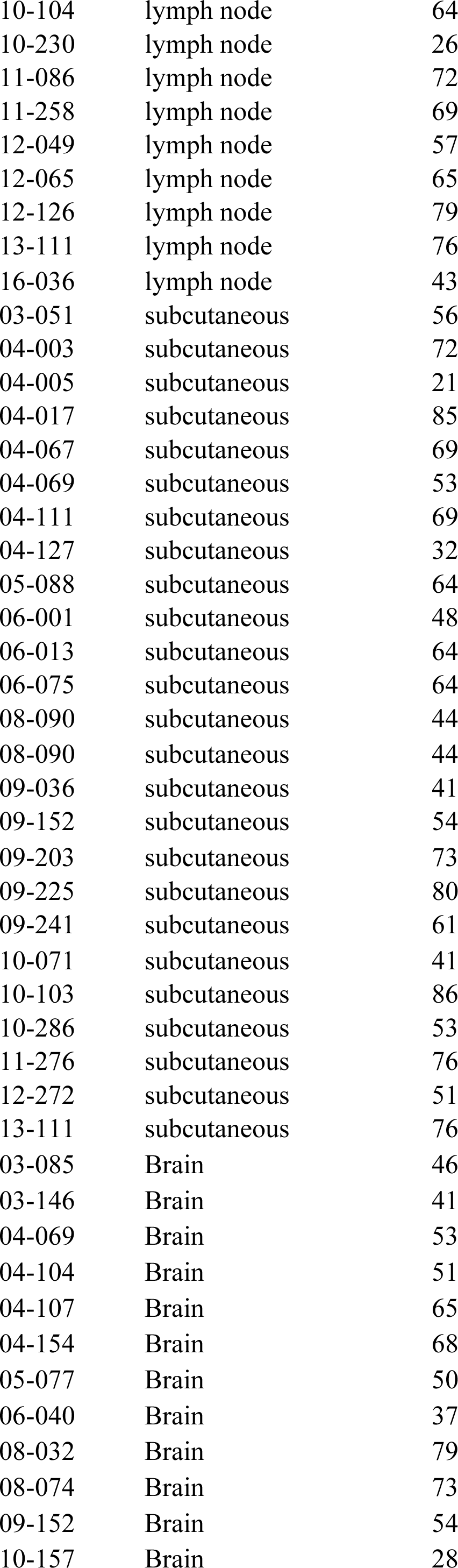
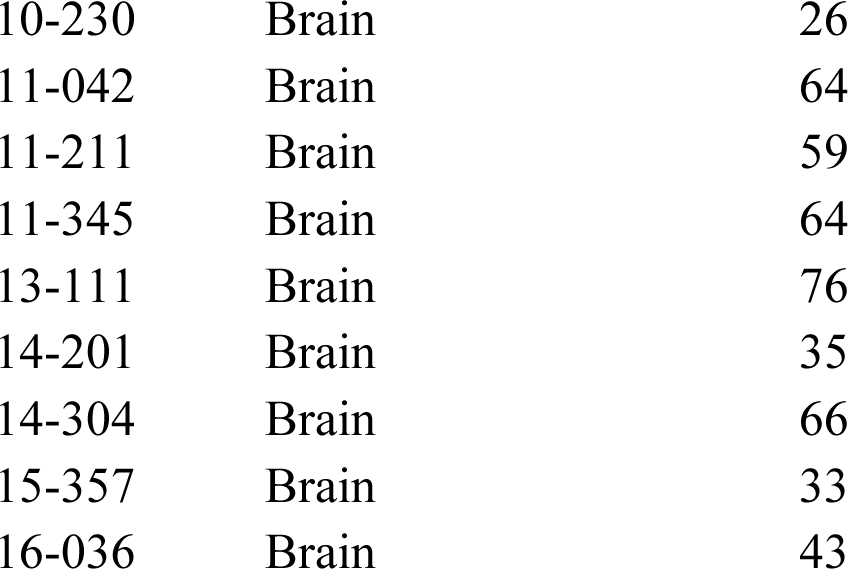
Human melanoma FFPE patient samples used in this study and their clinicopathological parameters.

**Table S3:**
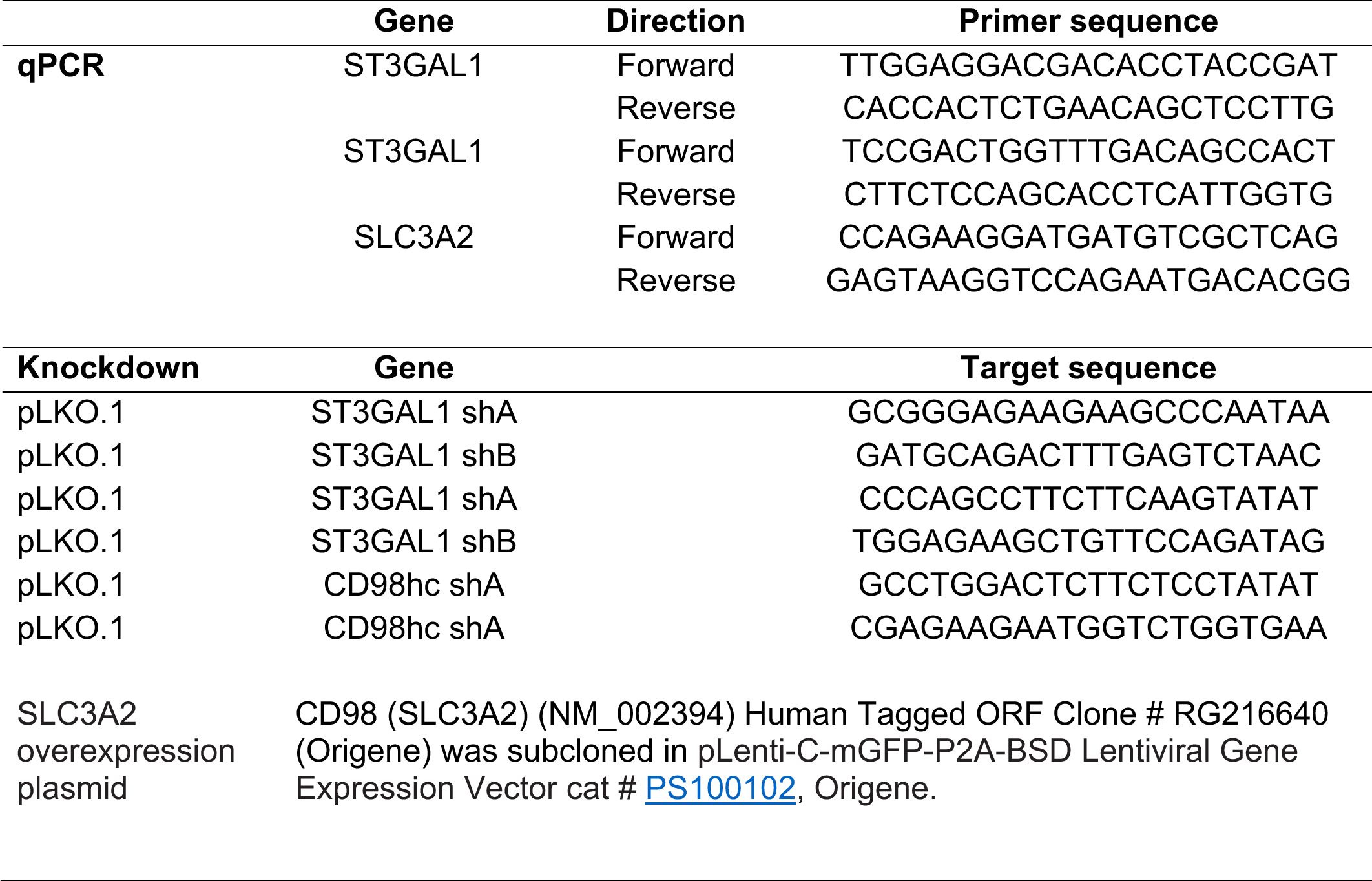
Primers, shRNA sequence, and constructs.

