## Supplementary table 3, https://www.origene.com/catalog/vectors/lentiviral-gene-expression-vectors/ps100102/plenti-c-mgfp-p2a-bsd-lentiviral-gene-expre for "Integrated *in vivo* functional screens and multi-omics analyses identify α-2,3-sialylation as essential for melanoma maintenance"

**Supplementary Material**

**Supplementary figures:**

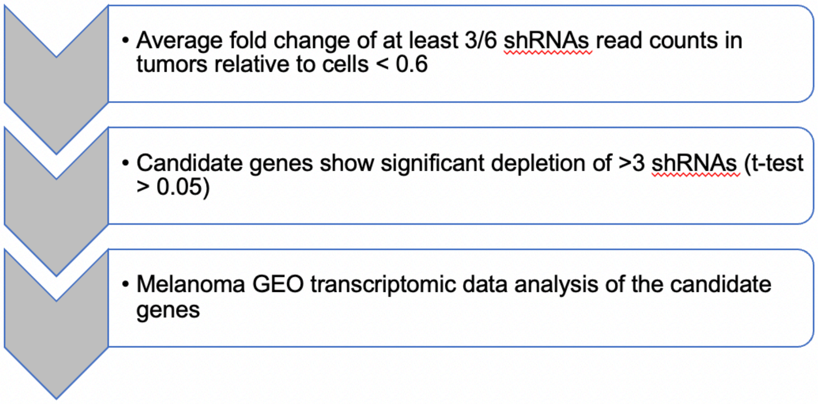


| Gene symbols | # of shRNAs (fold change < 0.6/ # of designed shRNAs |
|---|---|
| B3GAT1 | 5/6 |
| B3GNT4 | 4/6 |
| B3GNT5 | 4/6 |
| GALNT2 | 4/6 |
| GCNT2 | 4/6 |
| HS3ST6 | 4/6 |
| OGT | 6/6 |
| RFNG | 3/6 |
| RPN2 | 5/6 |
| ST3GAL1 | 3/6 |
| ST3GAL2 | 3/6 |
| ST3GAL5 | 3/6 |
| ST3GAL6 | 3/6 |
| ST6GALNAC2 | 4/6 |
| ST8SIA2 | 6/6 |
| UGT8 | 4/6 |
| XYLT1 | 3/5 |

**Fig S1:** Selection of glycogene candidates using *in vivo* shRNA functional screen (**A**) Stepwise filtering of depleted shRNA. (**B**) List of glycogenes for which # of shRNAs were consistently depleted.

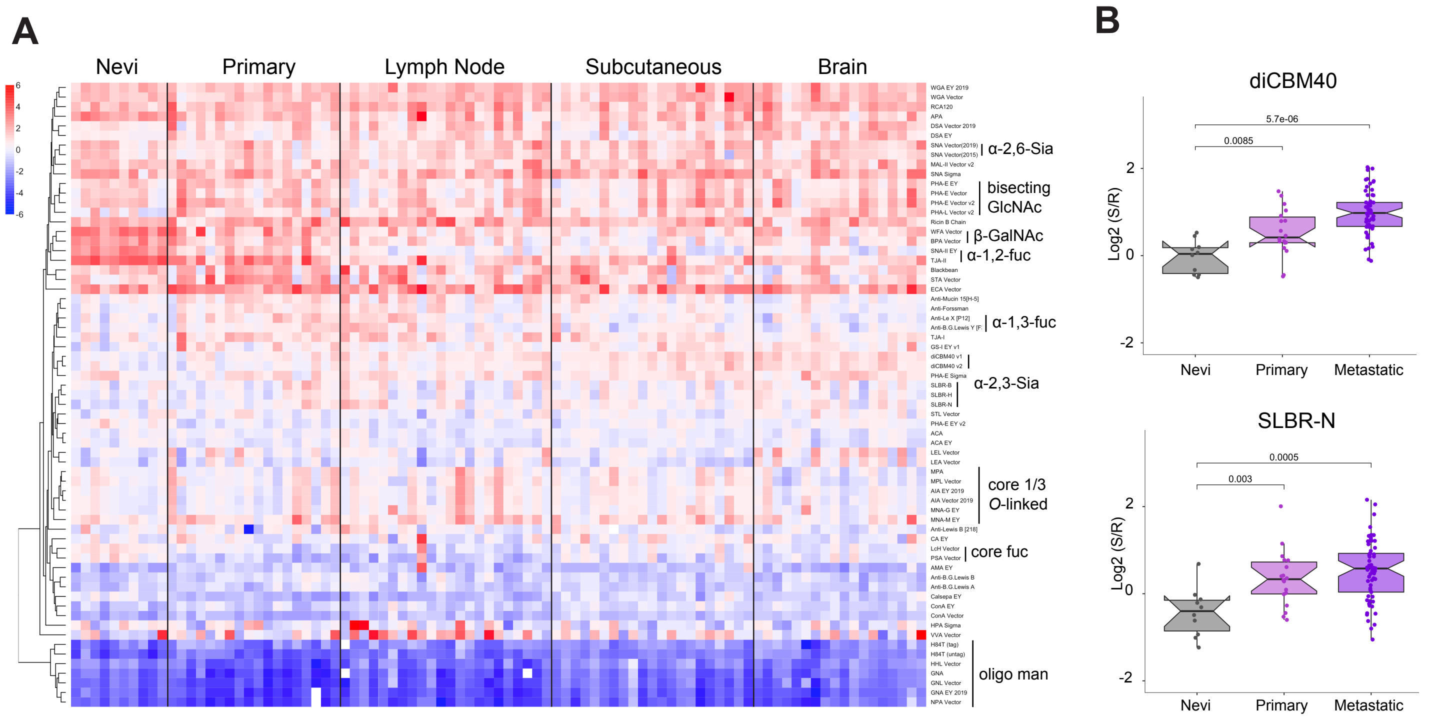


**Fig S2:** Glycome of melanoma differs from melanocytic nevi. (**A**) Glycomic profile of nevi (n = 10), primary melanoma (n = 18), lymph node metastasis (n = 22), subcutaneous metastasis (n = 21) brain metastasis (n = 18) by ratiometric lectin microarray data. All specimens were micro-dissected from formalin-fixed, paraffin-embedded tissues, and protein levels were determined by the BCA method. Equal amounts (5.0 µg by proteins) of Cy-5 labeled samples, and Cy-3 labeled references were analyzed by lectin microarray. A heatmap of lectin microarray data for all lectins is shown. Median normalized $\log_2$ ratios (Sample (S)/Reference(R)): red, $\log_2(S) > \log_2(R)$; blue, $\log_2(R) > \log_2(S)$. Lectins were hierarchically clustered using the Pearson correlation coefficient and average linkage analysis. Lectins bindings are highlighted to the right of the heatmap. (**B**) Boxplot analysis of **α**-2,3 sialosides probed by lectins diCBM40 and SLBR-N. Significance was determined using Student's t-test (two-tailed).

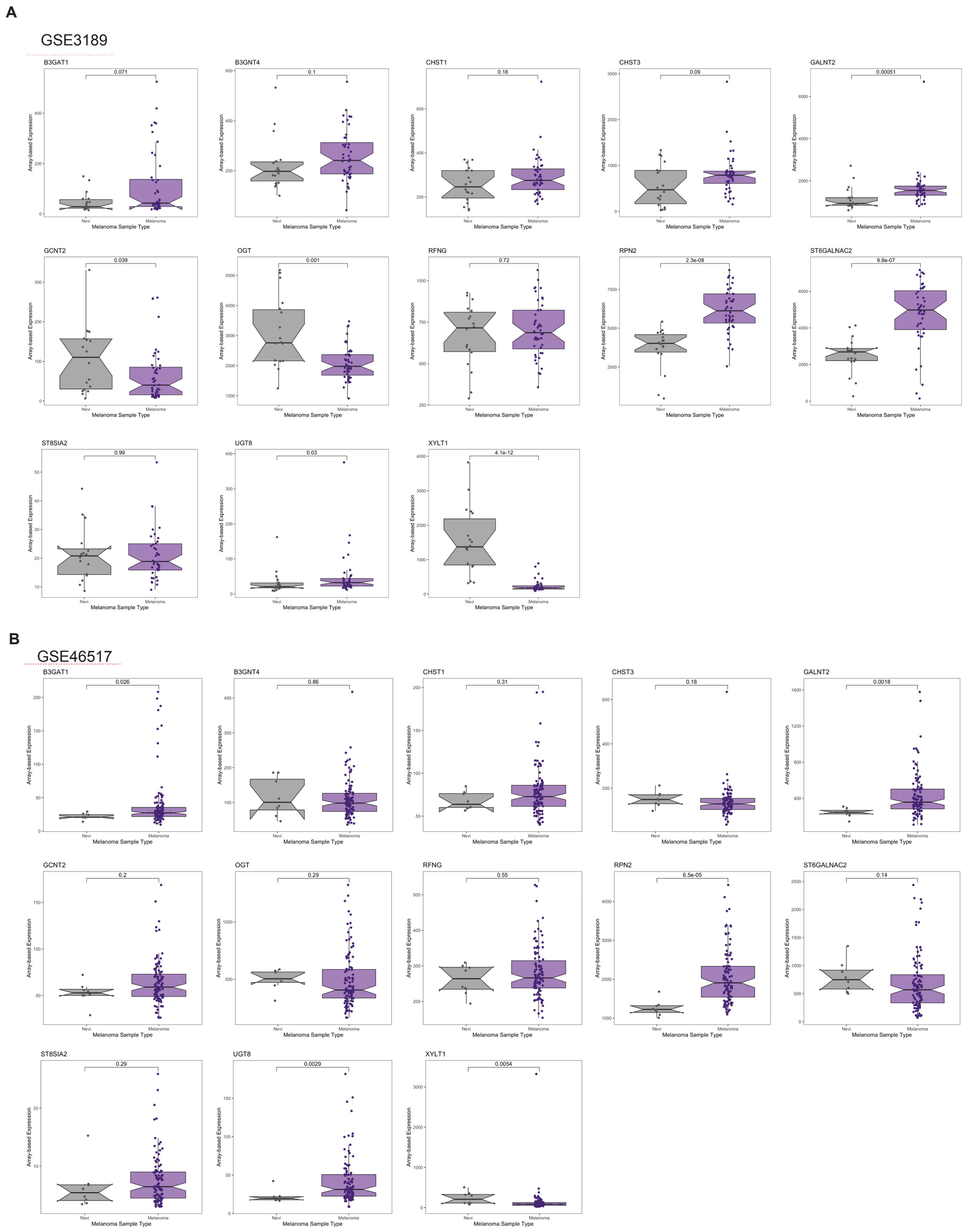
A
GSE3189
B3GAT1
0.071
B3GNT4
0.1
CHST1
0.18
CHST3
0.09
GALNT2
0.00051
GCNT2
0.039
OGT
0.001
RFNG
0.72
RPN2
2.3e-08
ST6GALNAC2
9.8e-07
ST8SIA2
0.99
UGT8
0.03
XYLT1
4.1e-12
B
GSE46517
B3GAT1
0.026
B3GNT4
0.86
CHST1
0.31
CHST3
0.18
GALNT2
0.0018
GCNT2
0.2
OGT
0.29
RFNG
0.55
RPN2
6.5e-05
ST6GALNAC2
0.14
ST8SIA2
0.29
UGT8
0.0029
XYLT1
0.0054
Array-based Expression
Nevi
Melanoma
Melanoma Sample Type

**C**

GSE12391

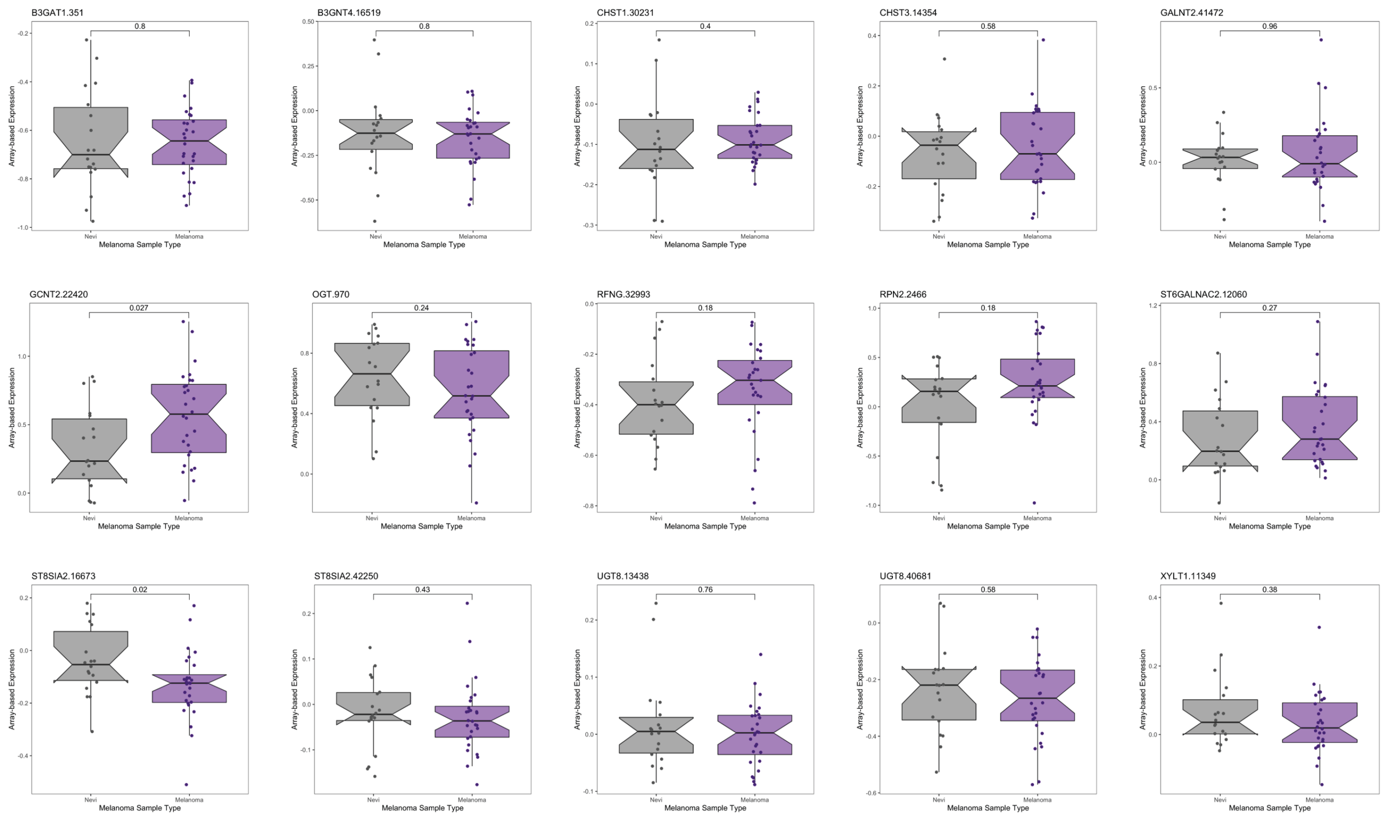


**Fig. S3: Expression of candidate glycogenes in melanoma relative to nevi.** Whisker plot illustrating mRNA expression level in melanoma samples compared to nevi in multiple datasets: **(A)** GSE3189(81), **(B)** GSE46517(82), **(C)** GSE12391 (83). Various glycogene candidates from *in vivo* functional library screening are presented in fig. S1B. Two-tailed unpaired t-test.

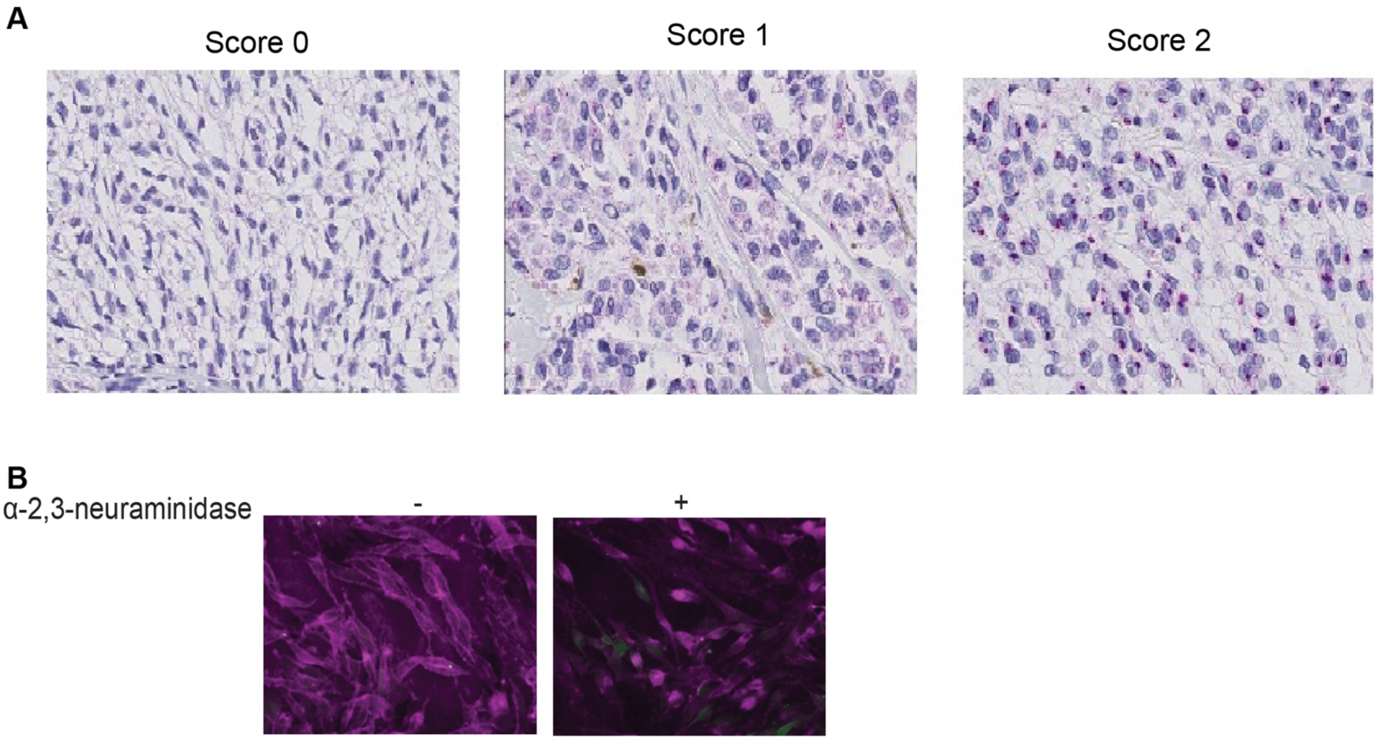


**Fig. S4: (A)** Representative ST3GAL1 IHC images depict IHC scoring categories. (**B**) diCBM40 fluorescence microscopy of 5B1 cells with or without α-2,3-neuraminidase (NEB, P0743) treatment. 5B1 cells were incubated with his tagged diCBM40 after neuraminidase treatment and visualized by 6x-His Tag Monoclonal Antibody (HIS.H8), Alexa Fluor™ 647.

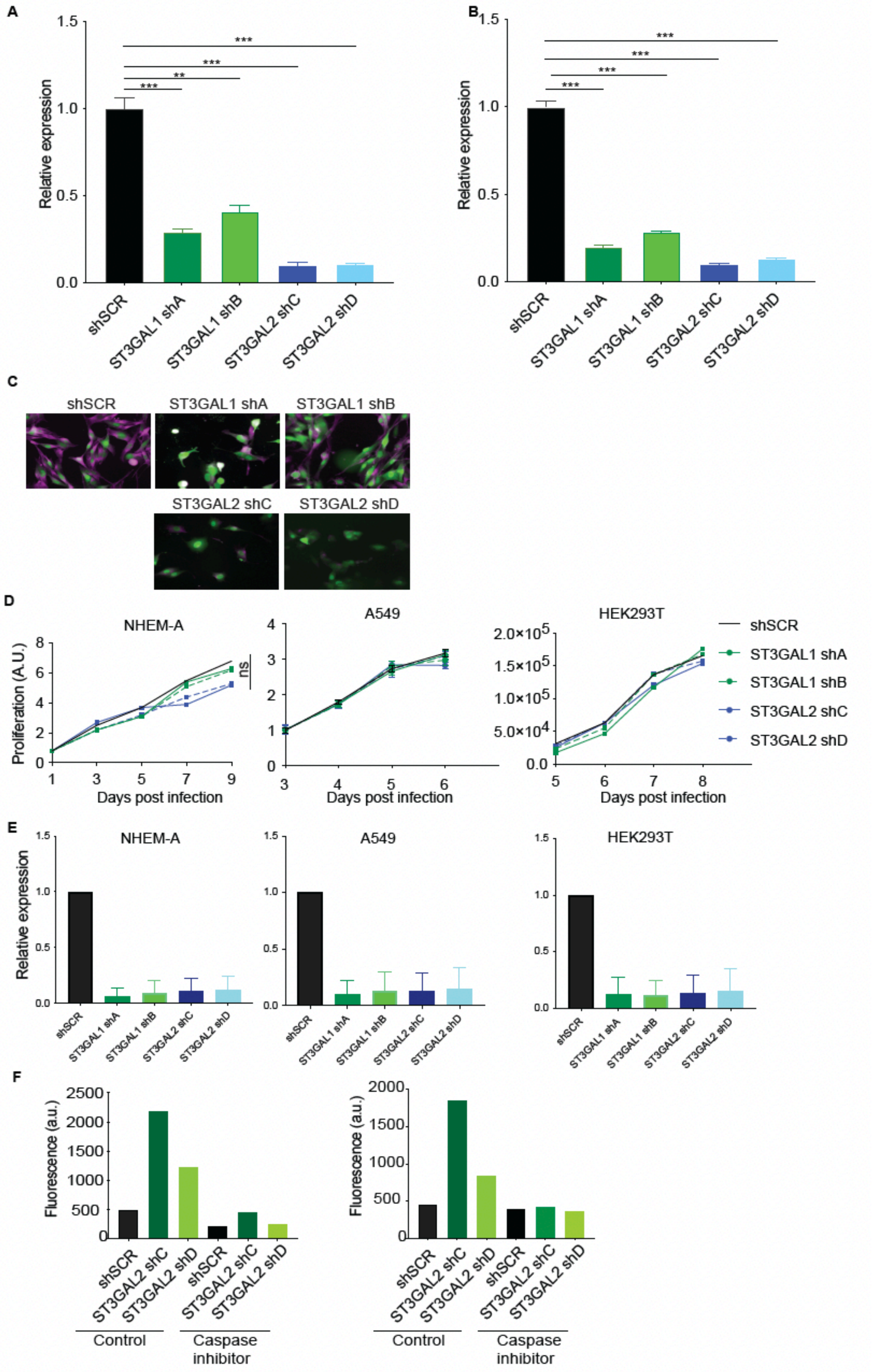


**Fig. S5: ST3GAL1 and ST3GAL2 are essential for melanoma proliferation *in vitro*.** (**A and B**) ST3GAL1 and ST3GAL2 transcript expression in 5B1 and 12-273BM cells stably expressing shRNA targeting ST3GAL1 (shA or shB) or ST3GAL2 (shC or shD) or shSCR were assessed by real-time qPCR. qPCR graph shows average relative expression normalized to *GAPDH*, three replicates per condition, two-tailed unpaired t-tests. qPCR data are representative of three independent experiments. (**C**) diCBM40 fluorescence microscopy of 5B1 cells transduced by ST3GAL1 (shA or shB) or ST3GAL2 (shC or shD) or shSCR. 5B1 cells were visualized by incubating with diCBM40 lectin followed by 6x-His Tag Monoclonal Antibody (HIS.H8), Alexa Fluor™ 647. (**D**) Relative growth curves and (**E**) real-time qPCR of NHEM-A, A549, and HEK293T cells stably transduced with non-targeting scrambled control shRNA (shSCR), *ST3GAL1* shRNAs (shA and shB) and *ST3GAL2* shRNAs (shC and shD). (**F**) FACS analysis of cleaved caspase-3 on melanoma cells, 5B1 (left) and 12-273BM (right) transduced with two independent shST3GAL2 (shC and shD) treated with caspase inhibitor or vehicle.

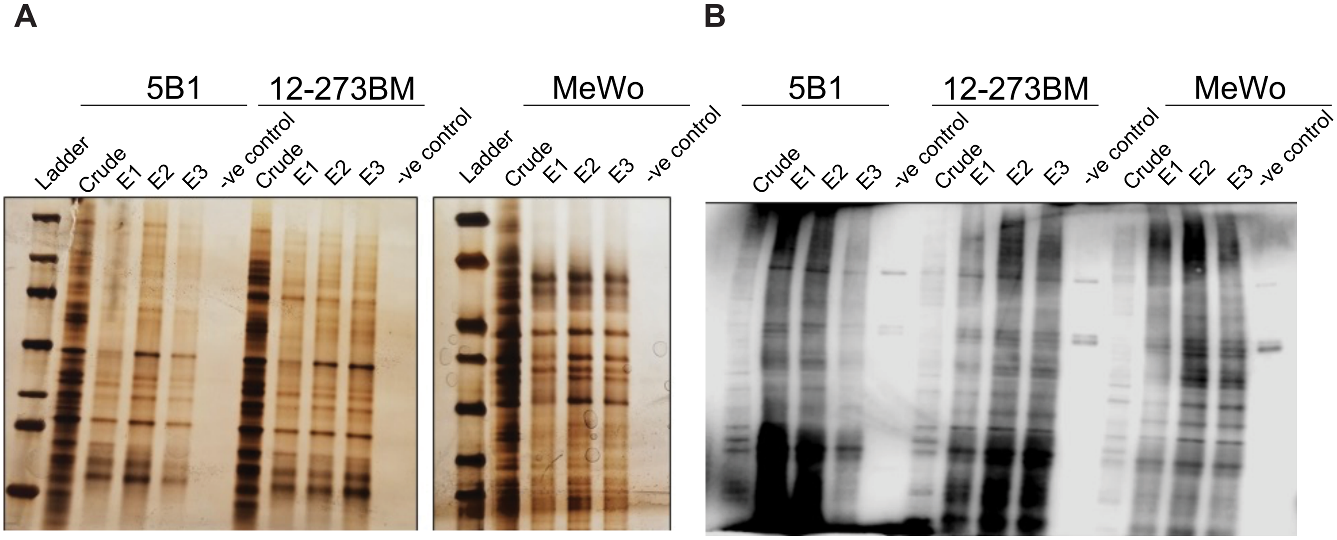


**Fig. S6: Identification of α-2,3-sialylated glycoproteins in melanoma.** Evidence of enrichment of α-2,3 sialylated proteins by **(A)** silver staining and **(B)** MAA lectin blot. An equal volume of MAA-enriched protein was loaded on the gel for silver stain or MAA lectin blot. The left lane shows a molecular weight marker (MW). Samples for each cell line were loaded in 3 biological replicates represented by E1, E2, and E3, along with negative control (without MAA lectin in the enrichment process).

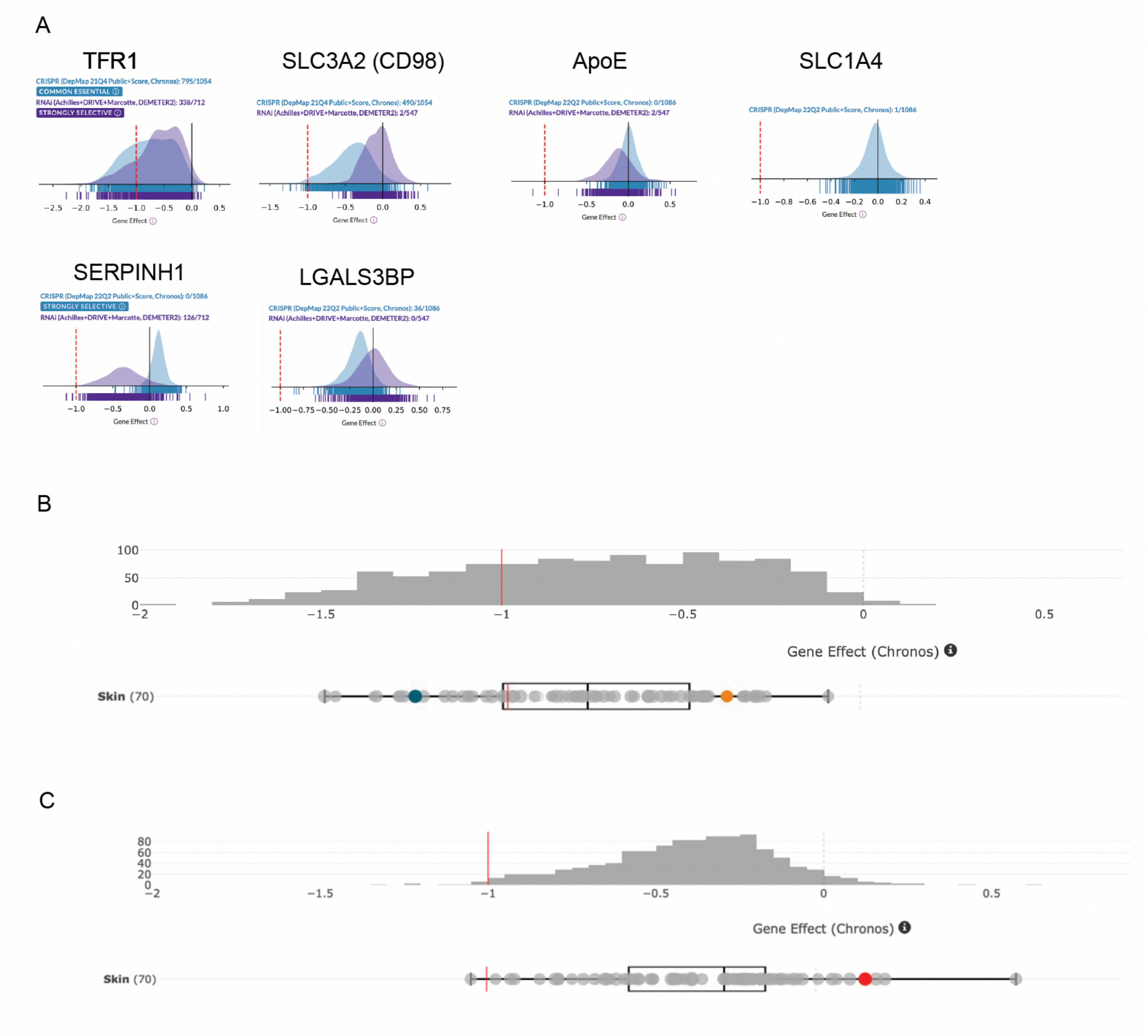


**Fig. S7: (A)** Association of sialylated proteins with an essentiality of genes: DepMap data mining of genes for various sialylated glycoproteins in cancer cells. **(B, C)** DepMap data mining of TFR1 and SLC3A2 in melanoma cell lines. Each dot represents a melanoma cell line. Gene effect (Chronos score) is shown. A lower Chronos score means a gene is more likely to be dependent in a given cell line. A score of -1 (red line) corresponds to the median of all common essential genes.

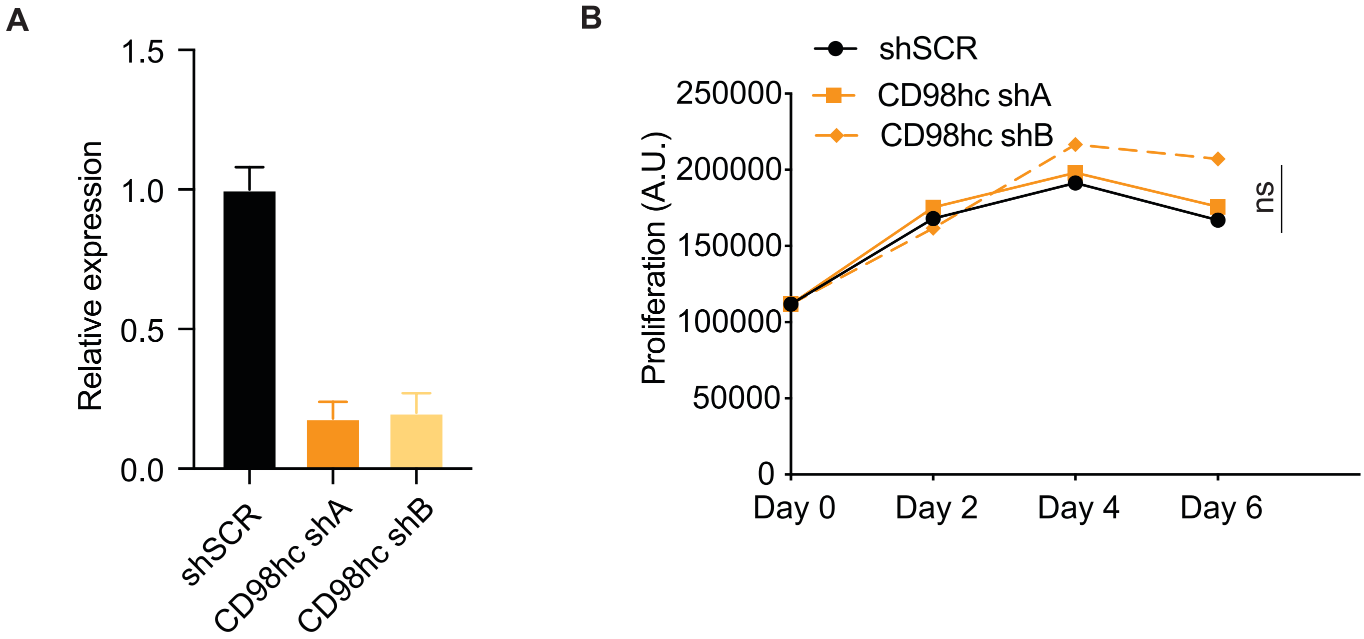


**Fig. S8: (A)** SLC3A2 transcript expression in HEK293T cells stably expressing shRNA targeting CD98hc (shA or shB) or shSCR was assessed by real-time qPCR. qPCR graph shows the average relative expression normalized to GAPDH. **(B)** The relative growth curves of HEK293T cells are stably transduced with non-targeting scrambled control shRNA (shSCR) or shRNA targeting CD98hc (shA or shB).

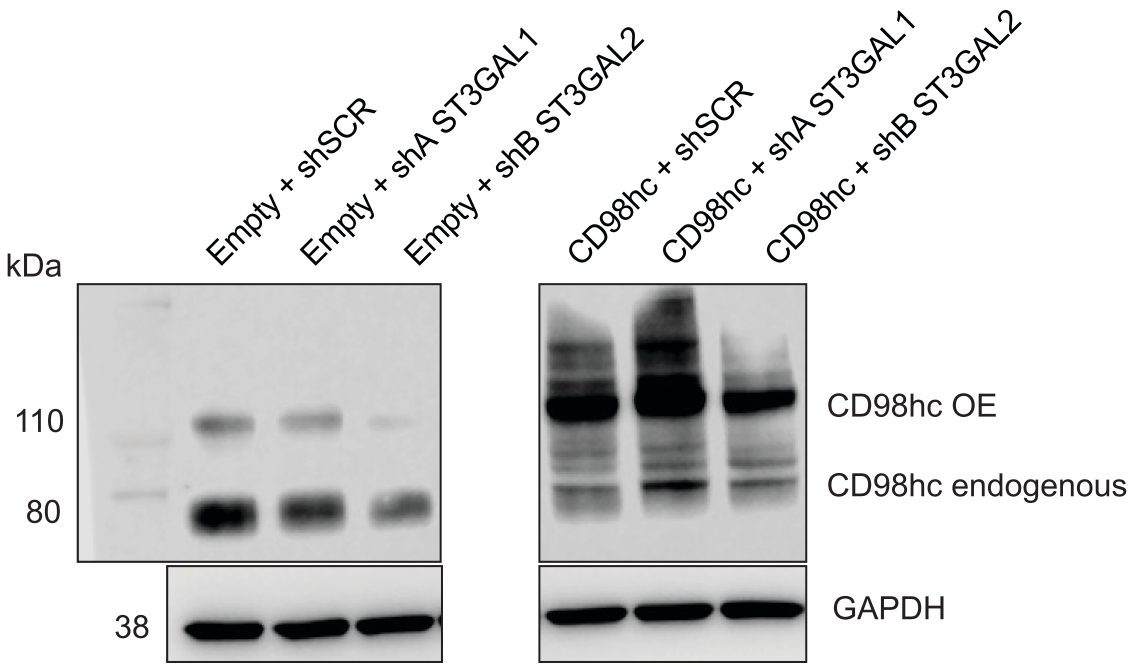


**Fig. S9:** Western blot for CD98hc on lysates from melanoma cells stably overexpressing CD98hc or empty vector and transduced with non-targeting control shSCR or shST3GAL1 or shST3GAL2.

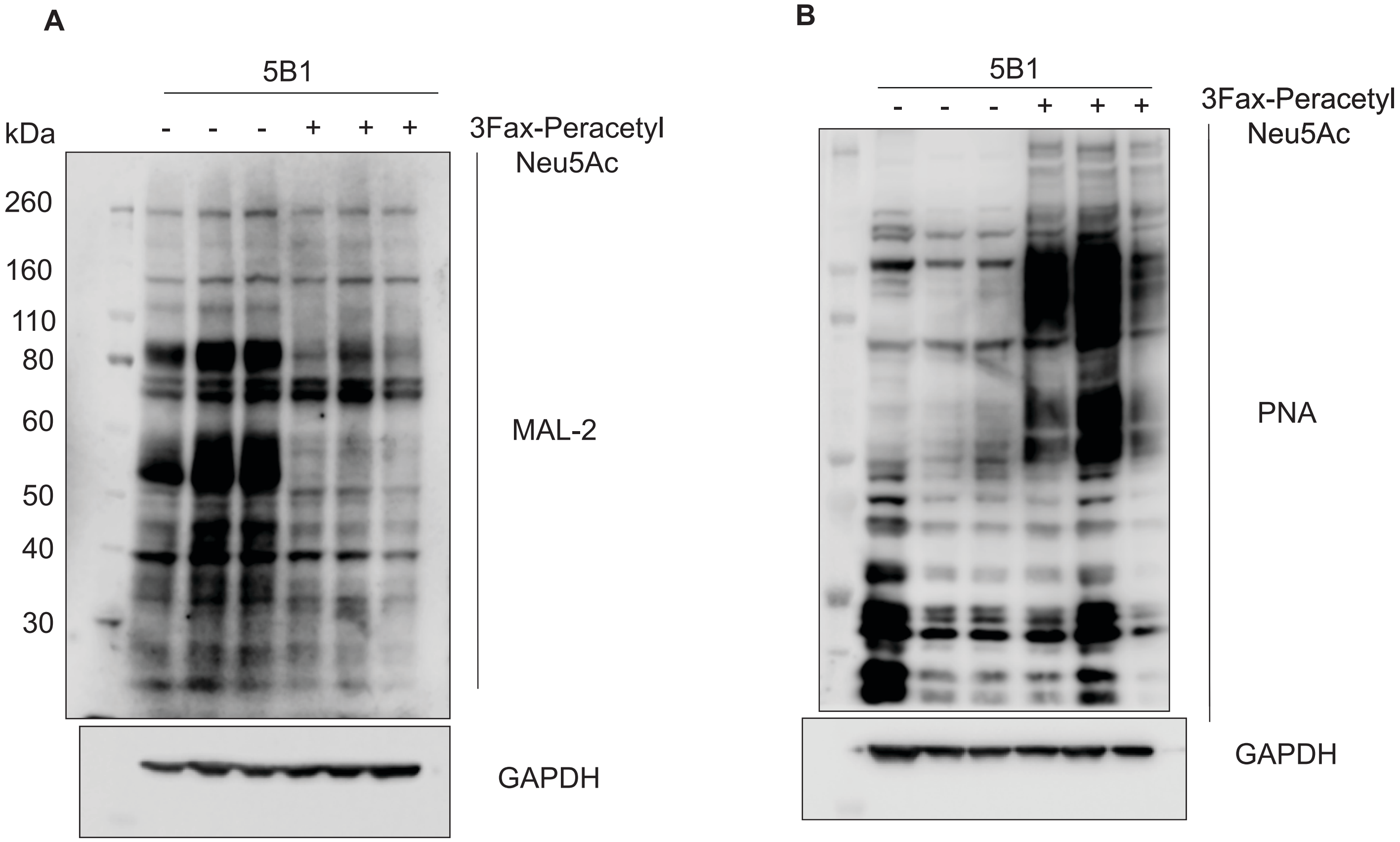


**Fig. S10:** Lectin blot analysis of 5B1 cells treated with sialyltransferase inhibitor 3Fax-peracetylNeu5Ac. **(A)** MAL-2 lectin blot **(B)** PNA lectin blot. GAPDH serves as a loading control.

**Supplementary Tables**

**Table S1:** Mass spectrometric analysis of MAA enriched proteins commonly present in the growth category of GO analysis in 5B1, MeWo cells, and 12-273BM STC.

| Accession# | Description | #PSM 12-273BM | | | #PSM 5B1_ | | | #PSM MeWo | | |
|---|---|---|---|---|---|---|---|---|---|---|
| **P50454** | Serpin Family H Member 1 (SERPINH1) | 58 | 58 | 42 | 28 | 42 | 33 | 29 | 33 | 34 |
| **P08195** | Isoform 2 of 4F2 cell-surface antigen heavy chain (SLC3A2, CD98hc) | 14 | 21 | 10 | 20 | 25 | 20 | 36 | 36 | 36 |
| **Q08380** | Galectin-3-binding protein (LGALS3BP) | 9 | 9 | 10 | 9 | 11 | 6 | 15 | 20 | 22 |
| **P43007** | Neutral amino acid transporter A (SLC1A4) | 17 | 15 | 12 | 5 | 7 | 8 | 13 | 16 | 8 |
| **P02786** | Transferrin receptor protein 1 (TFRC) | 16 | 16 | 12 | 6 | 9 | 4 | 6 | 7 | 4 |
| **P02649** | Apolipoprotein E (ApoE) | 15 | 14 | 7 | 3 | 12 | 6 | | | |

**Footnote:** The number of peptide spectral matches (PSMs) in each cell line.

**Table S2:** Human melanoma FFPE patient samples used in this study and their clinicopathological parameters.

| Pt. ID | Tissue type | Age at diagnosis |
|---|---|---|
| 04-066 | primary | 54 |
| 04-069 | primary | 53 |
| 04-116 | primary | 77 |
| 05-017 | primary | 75 |
| 06-002 | primary | 29 |
| 07-232 | primary | 71 |
| 09-036 | lymph node | 41 |
| 09-036 | subcutaneous | 41 |
| 09-152 | primary | 54 |
| 09-241 | primary | 61 |
| 10-040 | primary | 78 |
| 10-104 | primary | 64 |
| 10-175 | primary | 54 |
| 10-188 | primary | 60 |
| 10-217 | primary | 67 |
| 11-042 | primary | 64 |
| 11-054 | primary | 76 |
| 11-140 | primary | 80 |
| 11-345 | primary | 64 |
| 12-028 | primary | 66 |
| 12-040 | primary | 67 |
| 12-049 | primary | 57 |
| 12-065 | primary | 65 |
| 12-126 | primary | 79 |
| 12-148 | primary | 83 |
| 13-111 | subcutaneous | 76 |
| 02-017 | lymph node | 88 |
| 03-030 | lymph node | 76 |
| 04-069 | lymph node | 53 |
| 04-142 | lymph node | 74 |
| 04-157 | lymph node | 30 |
| 04-168 | lymph node | 25 |
| 05-017 | lymph node | 75 |
| 07-232 | lymph node | 71 |
| 08-145 | lymph node | 62 |
| 09-036 | lymph node | 41 |
| 09-085 | lymph node | 57 |
| 09-152 | lymph node | 54 |
| 09-241 | lymph node | 61 |
| 10-001 | lymph node | 87 |
| 10-040 | lymph node | 78 |

| | | |
|---|---|---|
| 10-104 | lymph node | 64 |
| 10-230 | lymph node | 26 |
| 11-086 | lymph node | 72 |
| 11-258 | lymph node | 69 |
| 12-049 | lymph node | 57 |
| 12-065 | lymph node | 65 |
| 12-126 | lymph node | 79 |
| 13-111 | lymph node | 76 |
| 16-036 | lymph node | 43 |
| 03-051 | subcutaneous | 56 |
| 04-003 | subcutaneous | 72 |
| 04-005 | subcutaneous | 21 |
| 04-017 | subcutaneous | 85 |
| 04-067 | subcutaneous | 69 |
| 04-069 | subcutaneous | 53 |
| 04-111 | subcutaneous | 69 |
| 04-127 | subcutaneous | 32 |
| 05-088 | subcutaneous | 64 |
| 06-001 | subcutaneous | 48 |
| 06-013 | subcutaneous | 64 |
| 06-075 | subcutaneous | 64 |
| 08-090 | subcutaneous | 44 |
| 08-090 | subcutaneous | 44 |
| 09-036 | subcutaneous | 41 |
| 09-152 | subcutaneous | 54 |
| 09-203 | subcutaneous | 73 |
| 09-225 | subcutaneous | 80 |
| 09-241 | subcutaneous | 61 |
| 10-071 | subcutaneous | 41 |
| 10-103 | subcutaneous | 86 |
| 10-286 | subcutaneous | 53 |
| 11-276 | subcutaneous | 76 |
| 12-272 | subcutaneous | 51 |
| 13-111 | subcutaneous | 76 |
| 03-085 | Brain | 46 |
| 03-146 | Brain | 41 |
| 04-069 | Brain | 53 |
| 04-104 | Brain | 51 |
| 04-107 | Brain | 65 |
| 04-154 | Brain | 68 |
| 05-077 | Brain | 50 |
| 06-040 | Brain | 37 |
| 08-032 | Brain | 79 |
| 08-074 | Brain | 73 |
| 09-152 | Brain | 54 |
| 10-157 | Brain | 28 |

| | | |
|---|---|---|
| 10-230 | Brain | 26 |
| 11-042 | Brain | 64 |
| 11-211 | Brain | 59 |
| 11-345 | Brain | 64 |
| 13-111 | Brain | 76 |
| 14-201 | Brain | 35 |
| 14-304 | Brain | 66 |
| 15-357 | Brain | 33 |
| 16-036 | Brain | 43 |

**Table S3:** Primers, shRNA sequence, and constructs

| | Gene | Direction | Primer sequence |
|---|---|---|---|
| **qPCR** | ST3GAL1 | Forward | TTGGAGGACGACACCTACCGAT |
| | | Reverse | CACCACTCTGAACAGCTCCTTG |
| | ST3GAL1 | Forward | TCCGACTGGTTTGACAGCCACT |
| | | Reverse | CTTCTCCAGCACCTCATTGGTG |
| | SLC3A2 | Forward | CCAGAAGGATGATGTCGCTCAG |
| | | Reverse | GAGTAAGGTCCAGAATGACACGG |

| Knockdown | Gene | Target sequence |
|---|---|---|
| pLKO.1 | ST3GAL1 shA | GCGGGAGAAGAAGCCCAATAA |
| pLKO.1 | ST3GAL1 shB | GATGCAGACTTTGAGTCTAAC |
| pLKO.1 | ST3GAL1 shA | CCCAGCCTTCTTCAAGTATAT |
| pLKO.1 | ST3GAL1 shB | TGGAGAAGCTGTTCCAGATAG |
| pLKO.1 | CD98hc shA | GCCTGGACTCTTCTCCTATAT |
| pLKO.1 | CD98hc shA | CGAGAAGAATGGTCTGGTGAA |
| SLC3A2 overexpression plasmid | CD98 (SLC3A2) (NM_002394) Human Tagged ORF Clone # RG216640 (Origene) was subcloned in pLenti-C-mGFP-P2A-BSD Lentiviral Gene Expression Vector cat # PS100102, Origene. | |
